# Highly accurate protein structure prediction-based virtual docking pipeline accelerating the identification of anti-schistosomal compounds

**DOI:** 10.1101/2025.06.09.658570

**Authors:** Wenjun Cheng, Mengjie Gu, Yuepeng Wang, Jing Wang, Shan Li, Gongwen Chen, Xu Chen, Oyetunde T. Oyeyemi, Yang Hong, Wei Hu, Jipeng Wang

**Author notes:** These authors contributed equally to this work (Wenjun Cheng, Mengjie Gu). Corresponding Authors (WH) (JW).

## Abstract

Schistosomiasis is a major neglected tropical disease that lacks an effective vaccine and faces increasing challenges from praziquantel resistance, underscoring the urgent need for novel therapeutics. Target-based drug discovery (TBDD) is a powerful strategy for drug development. In this study, we utilized AlphaFold to predict the structures of target proteins from *Schistosoma mansoni* and *Schistosoma japonicum*, followed by virtual molecular screening to identify potential inhibitors. Among 202 potential therapeutic targets, we identified 37 proteins with high-accuracy structural predictions suitable for molecular docking with 14,600 compounds. This screening yielded 268 candidate compounds, which were further evaluated *in vitro* for activity against both adult and juvenile *S. mansoni* and *S. japonicum*. Seven compounds exhibited strong anti-schistosomal activity, with HY-B2171A (Carubicin hydrochloride, CH) emerging as the most potent. CH was predicted to target the splicing factor U2AF65, and knockdown of its coding gene *Smp_019690* resulted in a phenotype similar to CH treatment. RNA sequencing revealed that both CH treatment and *Smp_019690* RNAi disrupted splicing events in the parasites. Further studies demonstrated that CH impairs parasite viability by inhibiting U2AF65 function in mRNA splicing regulation. By integrating RNAi-based target identification with structure-based virtual screening, alongside *in vitro* phenotypic and molecular analyses of compound-treated schistosomes, our study provides a comprehensive framework for anti-schistosomal drug discovery and identifies promising candidates for further preclinical development.

**Author Summary:** Schistosomiasis is a debilitating disease caused by parasitic worms and affects over 230 million people worldwide. For decades, control of the disease has relied on a single drug, praziquantel, which is less effective against juvenile parasites and increasingly threatened by the risk of drug resistance. To accelerate the discovery of new treatments, we integrated advanced computational tools with molecular validation. Using AlphaFold-predicted structures of 37 key schistosome proteins, we performed large-scale virtual screening to identify promising drug candidates. Over 200 compounds were prioritized for phenotypic screening on two major human-infective schistosome species, leading to the discovery of seven with potent anti-parasitic activity. Among them, Carubicin hydrochloride showed strong efficacy by targeting a critical splicing factor, U2AF65. Inhibiting this protein—either chemically or through gene silencing— disrupted parasite movement, suppressed cell proliferation, and altered splicing events. Our work highlights the power of integrating AI-based protein modeling with phenotypic and mechanistic validation to accelerate drug discovery for neglected tropical diseases like schistosomiasis.

## Introduction

Schistosomiasis, a zoonotic parasitic disease caused by infection with *Schistosoma spp*, is a significant public health concern. Globally, schistosomiasis has been reported in 78 countries, with 52 classified as endemic [1]. It is estimated that approximately 230 million individuals are infected worldwide, while 700 million people live at risk of infection [2]. As of 2021, at least 251.4 million people required preventive treatment [3]. The World Health Organization ranks schistosomiasis as the second most neglected tropical disease, surpassed only by malaria. Notably, the disease can infect both humans and a wide range of domestic animals, exacerbating its impact on human health and hindering economic development.

Schistosomiasis is closely linked to inadequate living conditions and poor sanitation. A comprehensive approach is emphasized by effective control strategies, including the improvement of sanitation facilities, promotion of health education, access to safe water supplies, and provision of adequate medical care [4]. In addition, the development of vaccines is considered a critical strategy for combating schistosomiasis. However, no effective vaccine has been developed to date. Another alternative approach focuses on the elimination of intermediate host snails. Historically, molluscicides were widely used for snail control prior to the 1970s, but their significant ecological impacts have limited their long-term application [5]. Despite these efforts, each of the aforementioned methods has inherent limitations, necessitating the exploration of alternative strategies. Increasing attention has been directed toward the discovery of anti-schistosomal drugs designed to interrupt the parasite’s life cycle and thereby reduce transmission. Currently, praziquantel (PZQ) is the only drug used for large-scale treatment and control of schistosomiasis. However, PZQ exhibits efficacy solely against adult worms and does not prevent reinfection. Moreover, its prolonged and repeated use has been associated with decreased sensitivity of parasites to the drug, leading to reduced therapeutic efficacy and the emergence of drug-resistance [6]. These challenges underscore the urgent need for the development of novel anti-schistosomal drugs to accelerate the eradication of this disease.

Current drug discovery approaches include phenotypic drug discovery (PDD) and target-based drug discovery (TBDD). Both of them have contributed to the screening of the anti-schistosomal compounds. For PDD, it searches for compounds that produce a desired effect on a cellular or organismal level, regardless of the exact molecular target. Abdulla et al. developed a partially automated screen workflow that interfaces schistosomula with microtiter plate-formatted compound libraries [7]. This workflow enabled high-throughput, phenotypic *in vitro* screening of 2,160 structurally diverse natural and synthetic compounds. The implementation of this method markedly improved the efficiency and throughput of drug discovery for schistosomiasis. For the TBDD strategy, it focuses on identifying compounds that bind to a specific known molecular target. Following the sequencing and annotation of the major human schistosome genomes, several putative drug targets have been identified and subjected to investigation [8]. Notable examples of these targets include *S. mansoni* thioredoxin glutathione reductase (*Sm*TGR) [9], polo-like kinase 1 (*Sm*PLK1) and tyrosine kinase [10], histone deacetylases (HDACs) and sirtuins [11, 12], cysteine protease cathepsin B1 (*Sm*CB1) [13], serotonin (5-HT) GPCR [14], and *S. japonicum* 3-oxoacyl-ACP reductase (OAR) [15]. Research efforts targeting these molecules have yielded several promising compounds with the potential to inhibit schistosome activity effectively. Recently, we performed a large-scale RNAi screening and identified around 200 potential druggable targets in *S. mansoni* [16], which provides essential resource for the TBDD against schistosomes.

The TBDD workflow begins with the identification and validation of drug targets, followed by simulating the binding interactions between candidate compounds and the target. A critical prerequisite for these processes is the determination of the target protein’s three-dimensional (3D) structure. However, elucidating protein structures remains one of the most formidable challenges in biology. Despite advances in techniques such as nuclear magnetic resonance (NMR) spectroscopy [17], X-ray crystallography [18], and cryogenic electron microscopy (Cryo-EM) [19], only around 200,000 protein structures have been resolved to date, representing less than 0.1% of all known proteins (https://www.rcsb.org/) [20]. Unlike human proteins, for which thousands of crystal or cryo-EM structures have been resolved, very few three-dimensional structures of *Schistosoma* proteins are available in public databases. This lack of structural data presents a major bottleneck for TBDD, which relies heavily on accurate protein models to simulate molecular interactions. To overcome this limitation, computational structure prediction has become a valuable alternative. In December 2020, AlphaFold2 (AF2), a machine learning-based protein structure prediction model developed by DeepMind, achieved a groundbreaking milestone by winning the 14th Critical Assessment of Structure Prediction competition. In 2021, DeepMind, in collaboration with the European Molecular Biology Laboratory, launched the AlphaFold Protein Structure Database, providing high-precision 3D protein structure predictions. This publicly accessible database now contains over 200 million predicted protein structures, encompassing nearly all known protein sequences, including the human proteome and the proteomes of many other organisms [20]. Among these predictions, approximately 35% exhibit experimental-level accuracy, while an additional 45% are considered highly reliable [20]. The database has significantly advanced fundamental research, aiding in the understanding of protein functions and disease mechanisms. Furthermore, it has become an indispensable resource for drug discovery, enzyme engineering, and biotechnology, offering unprecedented opportunities for innovation in these fields. Importantly, this includes predicted structural models for proteins in *S. mansoni* and *S. japonicum*, offering new possibilities for rational drug design against previously inaccessible targets.

In this study, we selected 202 potential therapeutic targets, primarily based on our previous large-scale RNAi screening on *S. mansoni* [16], along with seven newly identified genes from recent RNAi experiments on *S. mansoni* and *S. japonicum*. Using the highly accurate predicted structures of 37 target protein among these based on AlphaFold Protein Structure Database, we performed virtual docking against MCE active compound library and identified top 200 compound targeting each protein. We then prioritized 268 compounds and evaluated their inhibitory effect on both adult and juvenile worms of *S. mansoni* and *S. japonicum*. In total, we identified 7 compounds that showed high anti-schistosomal activity. Among them, HY-B2171A (Carubicin hydrochloride, CH), as a compound binding to the target protein splicing factor U2AF65 (Smp_019690) exhibited the most powerful efficacy. We further demonstrated that this compound affects the worm activity by inhibiting the function of the splicing factor U2AF65 in regulating the mRNA splicing. This study integrates high-accuracy predicted structures of potential targets with molecular docking, as well as *in vitro* phenotypic and molecular assessments of compound-treated schistosomes, thereby providing a robust approach that could significantly accelerate the drug discovery process for schistosomiasis.

## Results

### Virtual docking for potential druggable targets in schistosomes

To identify potential molecular inhibitors against schistosomes, we selected 202 protein targets essential for parasite viability, as determined by RNAi studies on *S. mansoni and S. japonicum*. This included 195 genes from our previous study [16] and 7 newly identified genes (4 genes from *S. mansoni* and 3 genes from *S. japonicum*) from recent screenings (S1 Fig). As shown in Fig 1, protein structure predictions for these targets were conducted using AlphaFold. After excluding low-quality models with an average per-residue model confidence score (pLDDT) < 70, 37 high-quality protein models were retained for virtual screening (S2 Table). These included 20 enzymes and 6 signal recognition particles. Hydrogen atoms were added to the protein models using Schrödinger Maestro 11.4, followed by energy optimization with the OPLS2005 force field. The Sitemap tool was used to predict the binding sites for each model, with the center of the docking box (20 Å × 20 Å × 20 Å) defined as the best binding site (Site1).

**Fig 1.**
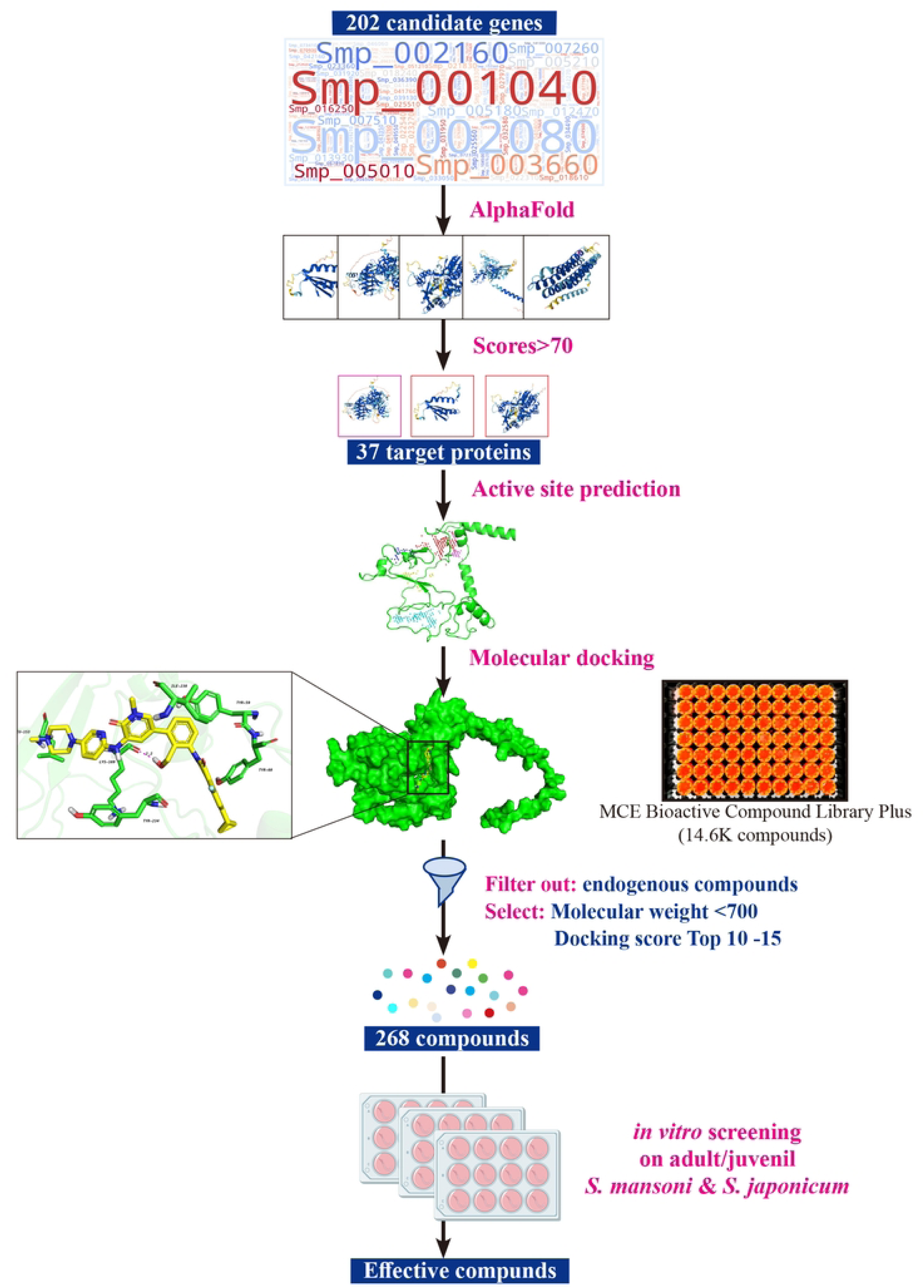
Workflow for the screening of putative anti-schistosomal compounds. This representation summarizes the inputs, outputs and filter criteria used in this study, leading to the selection of 268 compounds for *ex vivo* screening (see methodology for full details).

To identify potential active compounds targeting these proteins, we selected the MedChemExpress (MCE) Bioactive Compound Library Plus, which contains 14,600 compounds, for virtual docking (Fig 1). After adding hydrogen atoms and performing energy optimization on the 2D structures of the compounds (using the same procedure as for the protein models), 3D structures were generated. High-Throughput Virtual Screening (HTVS) was performed to identify primary compound candidates from the MCE library, with the top 10% subjected to Standard Precision (SP) docking. Subsequently, the top 10% of compounds from the SP results underwent a final round of docking using Extra Precision (XP) mode. The top 200 compounds with the highest docking scores for each target protein were selected (S3 Table). To prioritize the compounds, we excluded those with a molecular weight over 700 or endogenous compounds, and selected the top 10 -15 compounds with the highest docking scores for each protein target. In total, 268 compounds targeting 37 potential druggable proteins in *S. mansoni* and *S. japonicum* were identified from the MCE Bioactive Compound Library Plus (S4 Table).

### *In vitro* screening of effective compounds against adult *S. mansoni* and *S. japonicum*

During the virtual screening process, we observed a high degree of similarity between the target protein sequence of *S. mansoni* and its homolog in *S. japonicum* (S5 Table). Furthermore, protein structure predictions revealed a remarkable structural resemblance of the target protein in both species (S5 Table, S2 Fig). These findings suggest that the selected compounds may target the protein in both species, potentially inhibiting worm activity in *S. mansoni* and *S. japonicum*. Based on this, we conducted an extensive screening of compounds targeting both species.

To assess the effects of compounds on adult schistosomes, the viability of female and male worms was evaluated over seven days of culture in medium treated with 10 µM of the compounds. Key indicators of worm abnormalities included reduced attachment ability, swelling, shrinkage, darkened coloration, and abnormal motility. The activity of the cultured parasites was recorded daily, and the compound that caused 100% abnormality in worms on the final day of cultivation was considered effective, with PZQ used as a control.

For *S. mansoni* adult worms, a total of 26 effective compounds were identified (S6 Table), of which 13 exhibited pronounced effects, leading to worm mortality or a moribund state (Fig 2). These compounds targeted seven distinct protein targets, with phosphatidylserine decarboxylase proenzyme 1 (Smp_021830.3) being the most frequently targeted, followed by chitobiosyldiphosphodolichol alpha-mannosyltransferase (Smp_005010). Phenotypic analysis revealed significant activity for eight compounds (HY-100493, HY-15196, HY-112852, HY-12462, HY-12382, HY-B2171A, HY-12844, and HY-18018) after a single treatment, as observed on day 2 (D2) (S7 Table). Worms treated with these compounds exhibited key phenotypic changes, including dissociation of male-female pairs, failure to adhere to the bottom of the culture plates, reduced motility, and, in some cases, death (S7 Table). By D7, all highly effective compounds caused complete loss of adherence to the culture plate bottom, severely impaired viability, and states of death or near-death, characterized by intermittent, minimal movement restricted to the head or tail regions (S7 Table, Fig 2). Enlarged phenotypic images of worms treated with lethal compounds highlighted significant differences between the treatment and control groups (S3A Fig). In the control group, worms exhibited well-extended bodies and strong adherence to the substrate. In contrast, worms treated with either the effective compounds or PZQ displayed darkened coloration, shortened bodies, and, in some instances, curling into a circular posture. Additionally, worms in some treatment groups showed varying degrees of epidermal erosion, with blurred surface structures indicative of dissolution-like damage.

**Fig 2.**
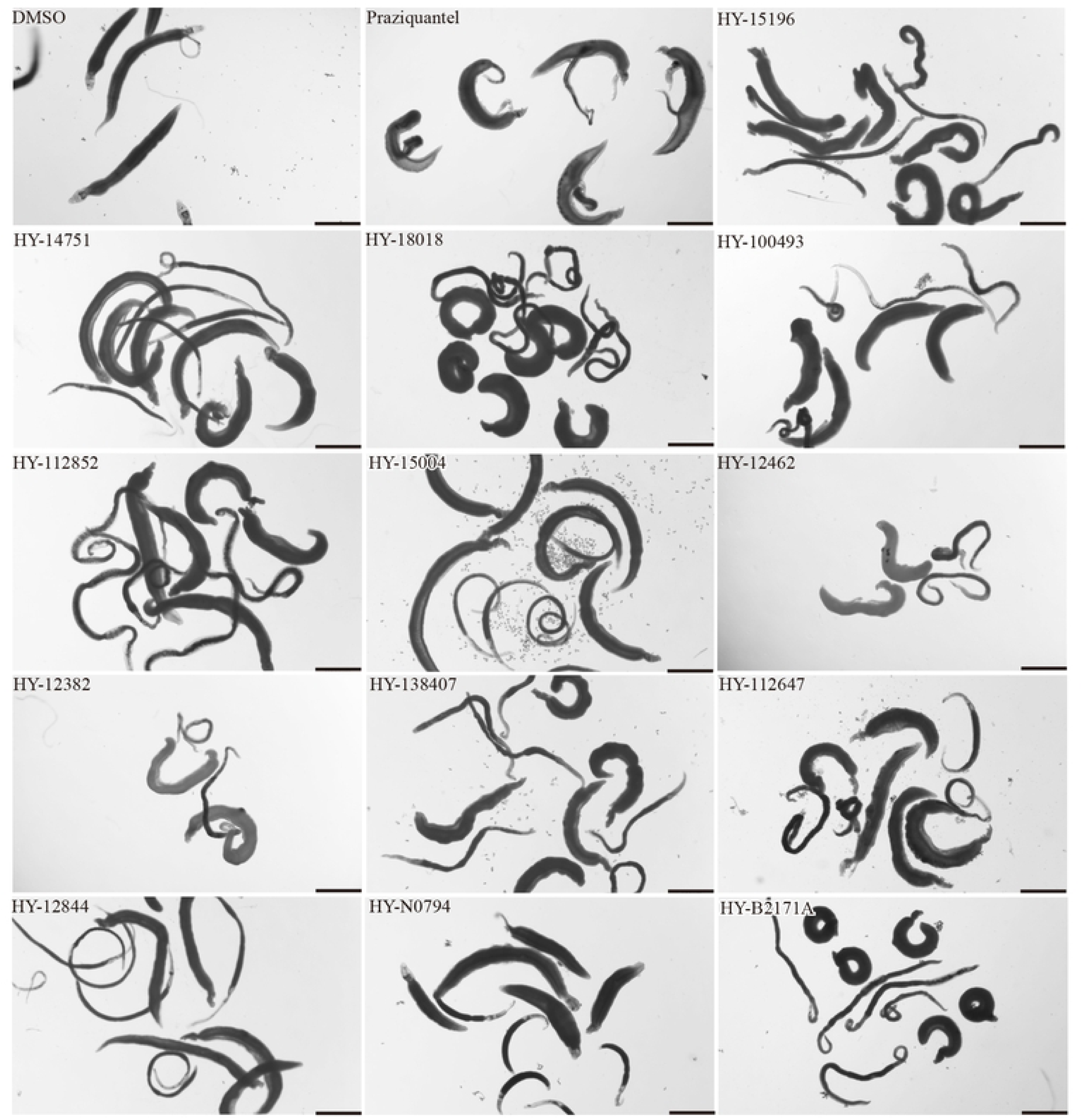
Phenotypes of adult *S. mansoni* treated with high-efficiency compounds. The figure depicts the phenotypic changes in *S. mansoni* after a 7-day co-incubation with thirteen high-efficiency compounds at a concentration of 10 μM. Negative control group: DMSO (0.1% DMSO, at the same DMSO concentration as in the experimental group.); Positive control group: Praziquantel (1 µM in 0.1% DMSO). Representative images from 3 biological replicates. Scale bars, 1000 μm.

To further evaluate their efficacy, these potent compounds were subjected to secondary screening at a lower concentration (1 µM), resulting in the identification of two compounds with significant effects (Table 1, S8 Table).

**Table 1.**
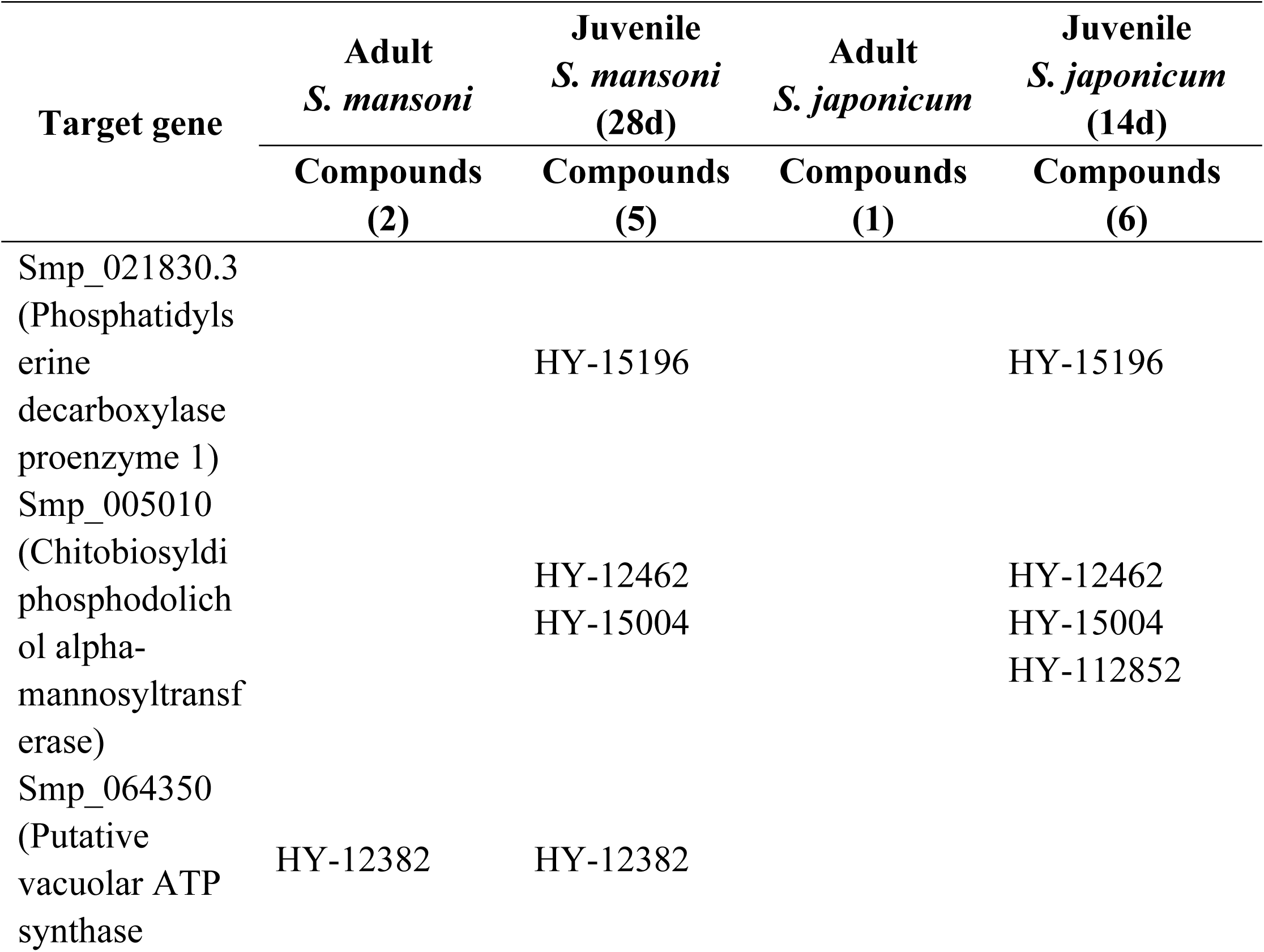
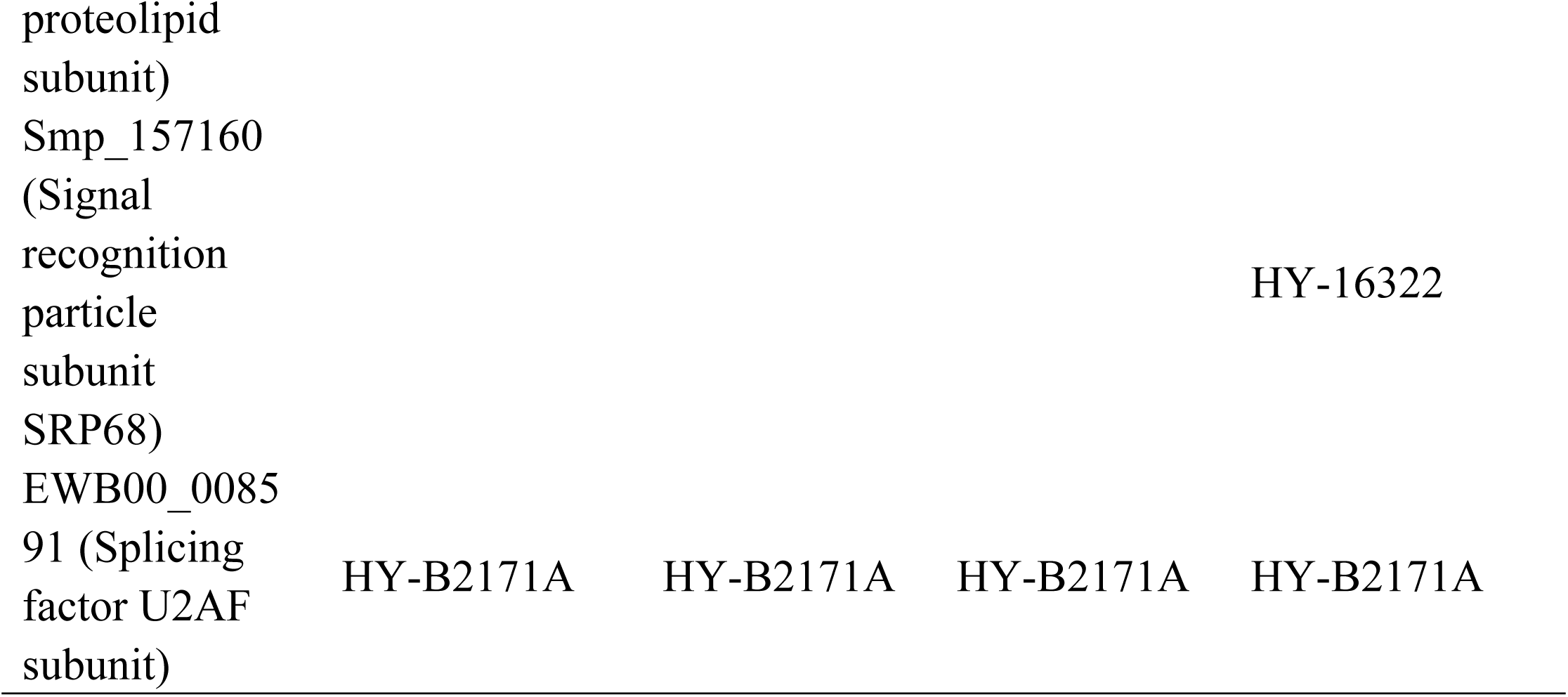
Compounds exhibiting potent inhibitory activity against *S. mansoni* and *S. japonicum* at 1 µM.

Similarly, an initial screening at 10 µM was conducted on *S. japonicum* adult worms, resulting in the identification of 23 effective compounds, 11 of which demonstrated high efficacy (S6 Table, Fig 3). The most frequently targeted proteins were consistent with those identified in *S. mansoni*, namely phosphatidylserine decarboxylase proenzyme 1 (Smp_021830.3) and chitobiosyldiphosphodolichol alpha-mannosyltransferase (Smp_005010). The detailed results of phenotypic recordings were presented in S9 Table. High-resolution images revealed the damages on the parasites (S3B Fig). Unlike the control group, worms in the PZQ group were dead, displaying darker coloration and a curled, ring-like posture. Additionally, worms in the treatment groups exhibited near-death characteristics, including darkened epidermal coloration, varying degrees of erosion and damage, and swelling. At a lower concentration of 1 µM, one compound (HY-B2171A) exhibited a significant effect (Table 1, S8 Table), with its predicted target identified as splicing factor U2AF subunit (EWB00_008591).

**Fig 3.**
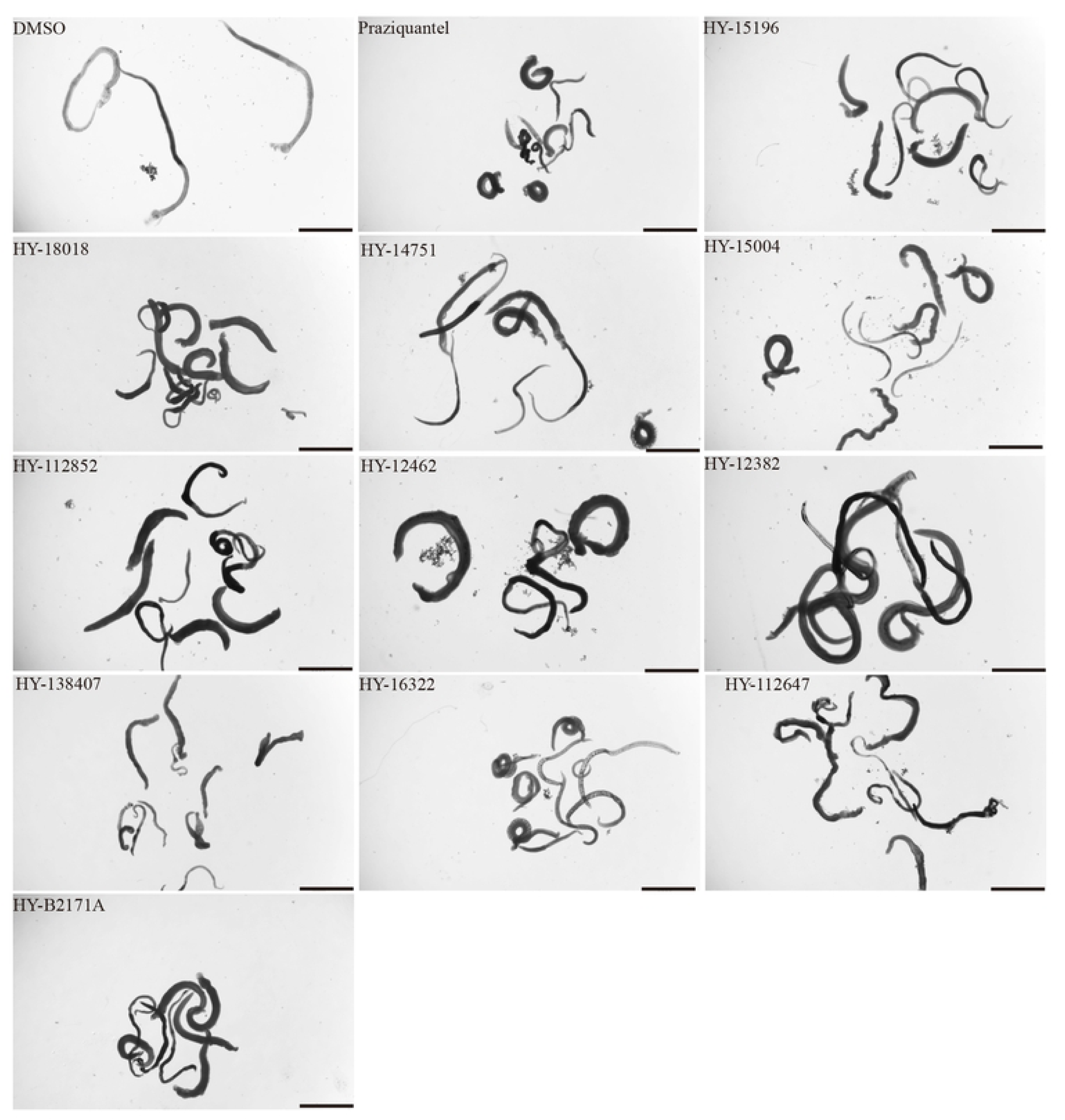
Phenotypes of adult *S. japonicum* treated with high-efficiency compounds. Bright field images of representative compound-induced phenotypes observed following 7-day co-incubation at a concentration of 10 μM. Negative control group: DMSO (0.1% DMSO, at the same DMSO concentration as in the experimental group.); Positive control group: Praziquantel (1 µM in 0.1% DMSO). Representative images from 3 biological replicates. Scale bars, 2000 μm.

### *In vitro* screening of effective compounds against juvenile *S. mansoni* and *S. japonicum*

Next, *in vitro* screening of compounds against juvenile worms was also performed. Compounds were deemed effective when all worms exhibited loss of attachment and reduced mobility. Those causing 100% mortality or moribundity were considered strongly active. Phenotypes for effective compounds were recorded in S10 and S11 Tables.

For juvenile *S. mansoni*, 29 compounds were identified with activity, 14 of which exhibited pronounced potency (S6 Table), inducing 100% mortality or moribund states (Fig 4). These most active compounds targeted seven distinct protein targets (S6 Table). Subsequently screening at a reduced concentration of 1 µM revealed five compounds with significant effects (Table 1, S8 Table). These five compounds targeted four distinct proteins, among these, two compounds—HY-12382, and HY-B2171A—were identical to those effective against adult worms (Table 1).

**Fig 4.**
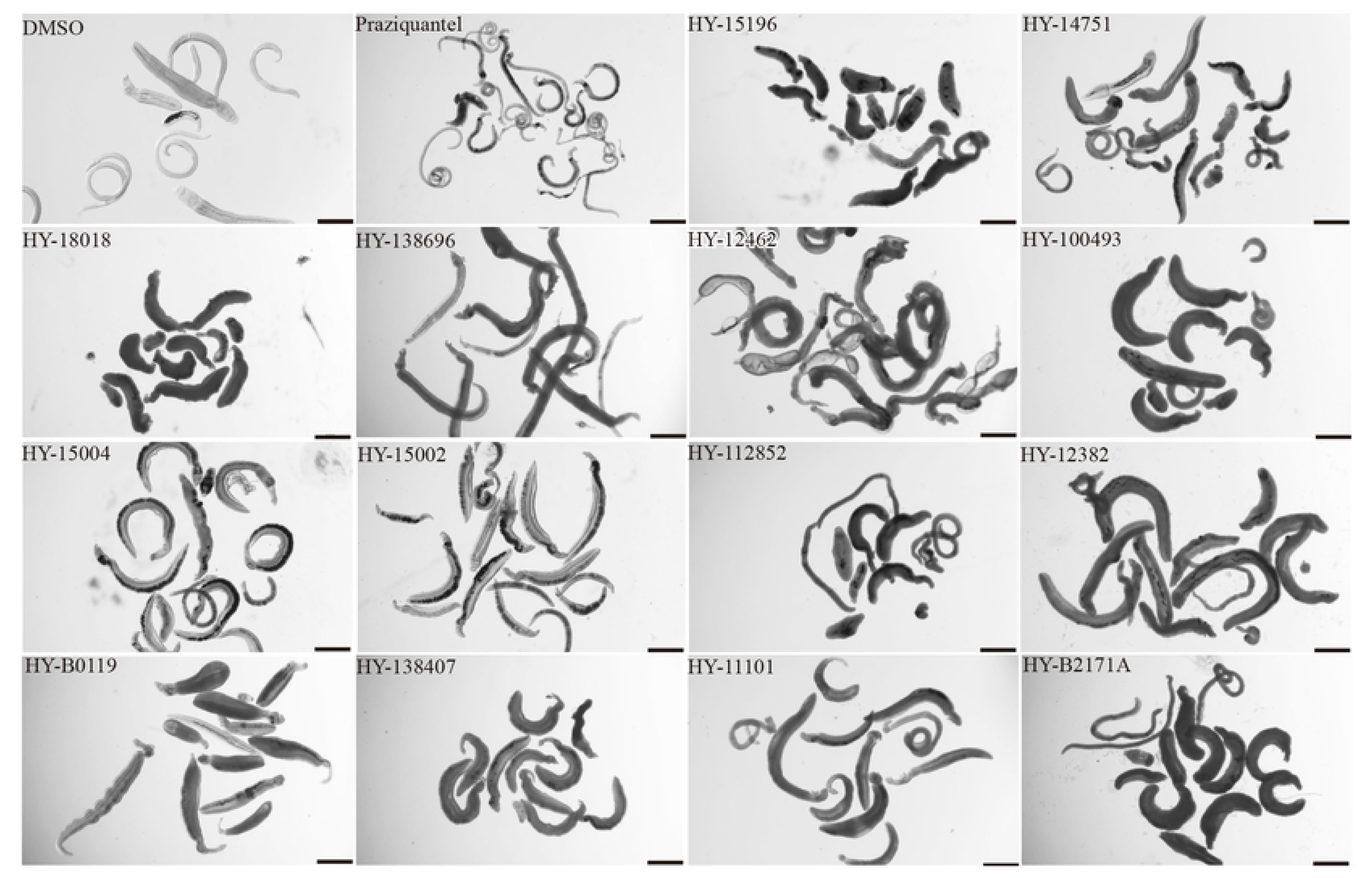
Phenotypes of juvenile *S. mansoni* treated with potent compounds. Fourteen compounds demonstrated superior inhibitory effects, causing 100% of the worms to lose attachment and exhibit reduced motility after a 7-day co-incubation at a concentration of 10 μM. 0.1% DMSO as negative control; 1 µM Praziquantel in 0.1% DMSO as positive control. Representative images from 3 biological replicates. Scale bars, 500 μm.

For *S. japonicum* juvenile, 29 effective compounds were identified, of which 16 demonstrated high efficacy (S6 Table, Fig 5). These strong inhibitors targeted seven distinct proteins, with one target differing from those identified in *S. mansoni*. This unique target (Smp_157160), corresponding to compound HY-16322, exhibited higher efficacy in *S. japonicum* than in *S. mansoni*. When the screening concentration was reduced to 1 µM, only six compounds remained effective (Table 1, S8 Table). The corresponding targets were Smp_021830.3, Smp_005010, Smp_157160, and EWB00_008591. Among these, Smp_157160 was unique to *S. japonicum*, while the other three targets were consistent with those identified in *S. mansoni*.

**Fig 5.**
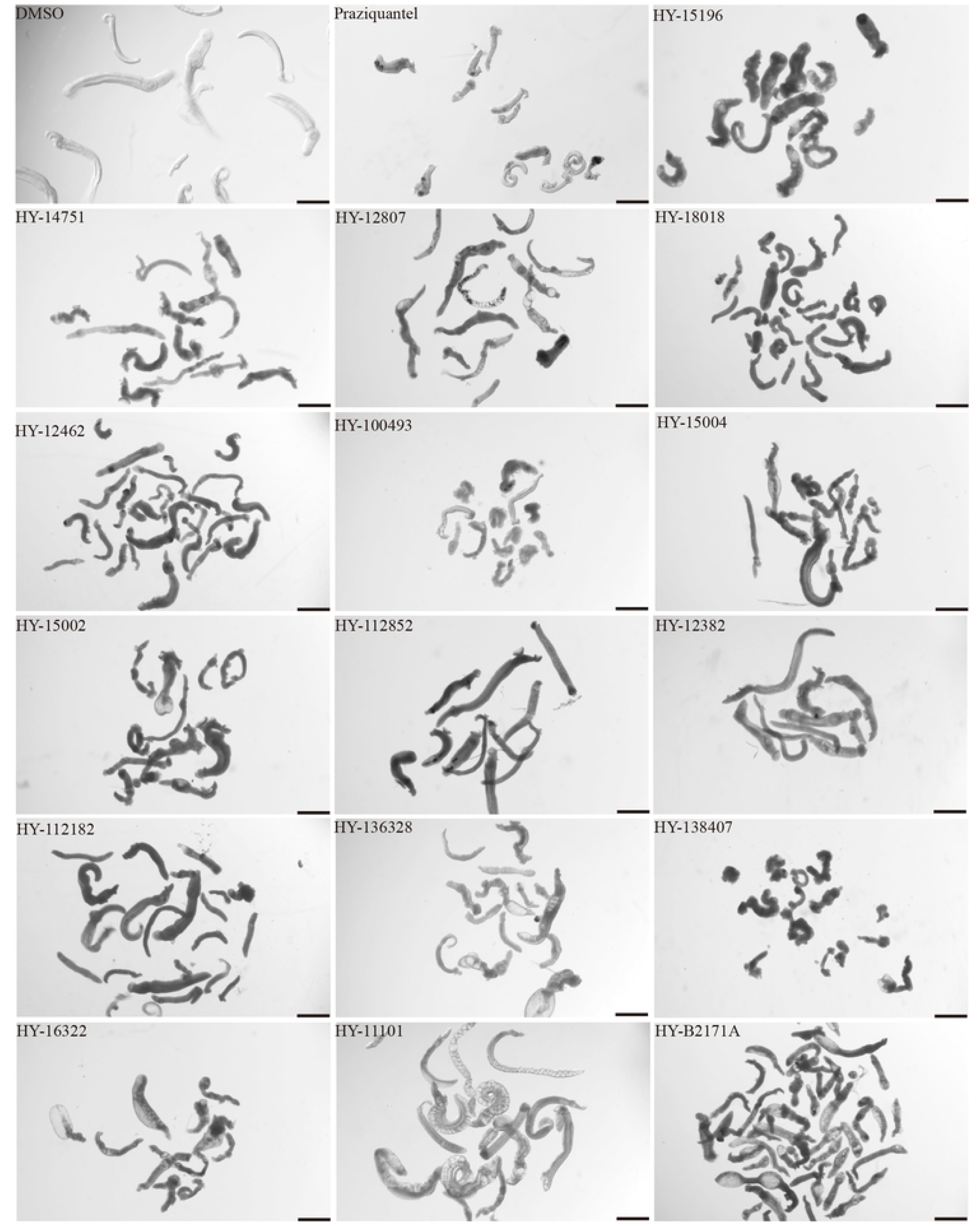
Phenotypes of juvenile *S. japonicum* treated with high-efficiency compounds. Bright-field images showcasing the phenotypes induced by sixteen representative compounds after a 7-day co-incubation at a concentration of 10 μM. 0.1% DMSO as negative control; 1 µM Praziquantel in 0.1% DMSO as positive control. Representative images from 3 biological replicates. Scale bars, 500 μm.

### Compounds demonstrating strong efficacy against both adult and juvenile schistosomes in two species

As previously described, at a concentration of 1 µM, seven compounds showed high anti-schistosomal activity, whose docking efficiency with targets were displayed in Fig 6A. In detail, two compounds demonstrated activity against *S. mansoni* adult worms (Fig 6B), whereas only one compound exhibited efficacy against *S. japonicum* adult worms (Fig 6C). For juvenile worms, four compounds were effective against both species; however, compound HY-12382 showed activity exclusively against *S. mansoni* juvenile (Fig 6D), while HY-112852 and HY-16322 were specifically effective against *S. japonicum* juvenile (Fig 6E). Notably, compound HY-B2171A (CH) exhibited broad-spectrum efficacy across all four types of schistosomes, demonstrating the most significant activity.

**Fig 6.**
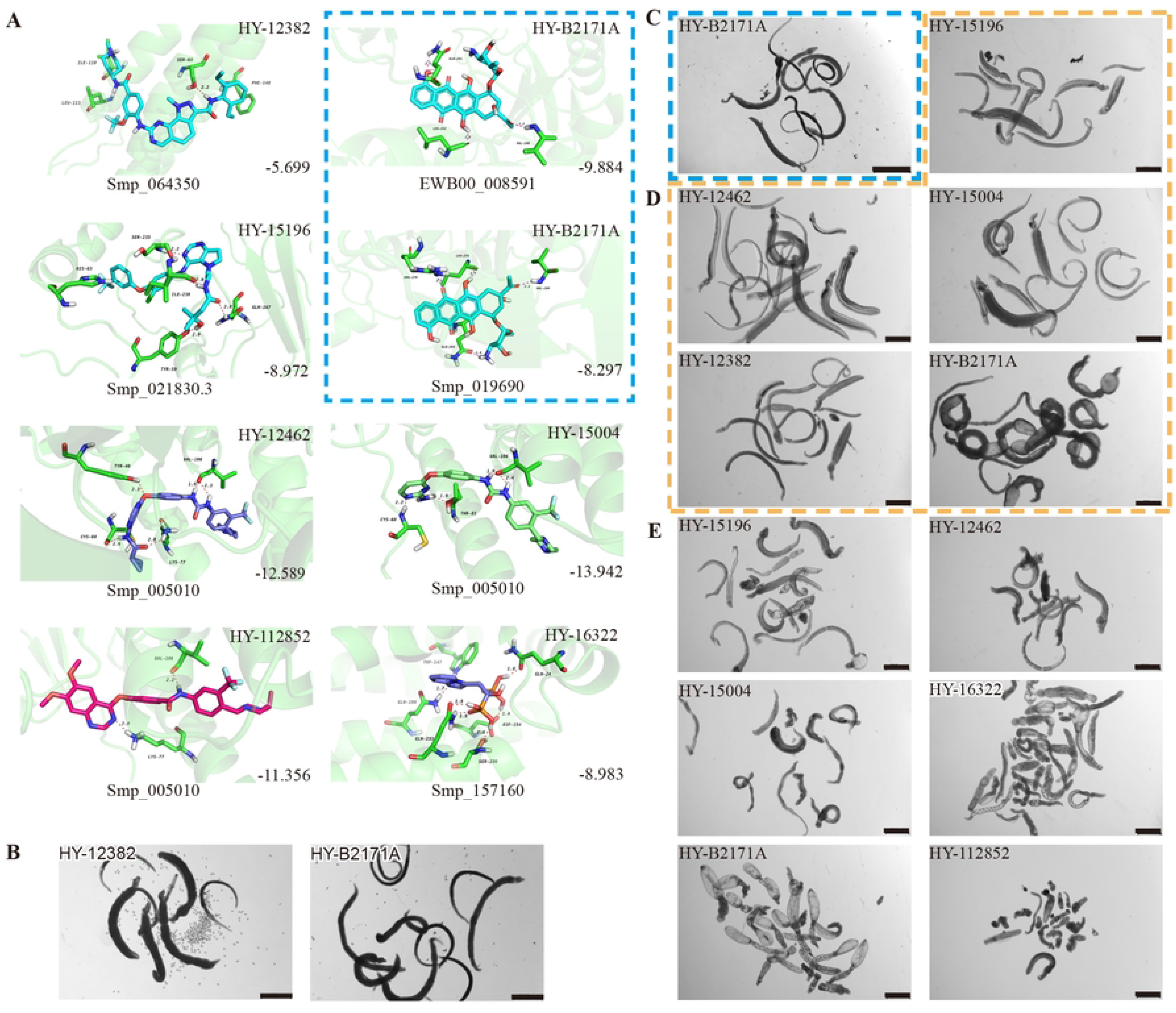
Molecular Docking of effective compounds with their targets and effects on schistosomes. (A) Molecular docking diagram of active compounds with the target. The C backbone of the target protein is depicted in green, nitrogen atoms (N) in blue, oxygen atoms (O) in red, and hydrogen atoms (H) in white. The compounds are shown as sticks with the following color codes: dark blue for HY-12462 and HY-16322, magenta for HY-112852, light green for HY-15004, and light blue for HY-15196, HY-12382, and HY-B2171A. π-π interactions are indicated by green dashed lines, and hydrogen bonds are shown as purple dashed lines, with longer hydrogen bond lengths reflecting weaker interactions. Light blue dashed box: molecular docking of compound CH with binding site of *Sj*U2AF65 (EWB00_008591) and *Sm*U2AF65 protein (Smp_019690). (B–E) Bright-field images showing representative compound-induced phenotypes following 7-day co-incubation with 1 μM compounds. (B) Adult *S. mansoni*; (C) Adult *S. japonicum*; (D) Juvenile *S. mansoni*; (E) Juvenile *S. japonicum*. Scale bars: B, 1000 μm; C, 2000 μm; D–E, 500 μm. Representative images from three biological replicates.

### CH is a potent anti-schistosomal compound

Based on previous results, nine compounds demonstrating significant efficacy against both adult schistosomes and juvenile were identified for *S. mansoni* and *S. japonicum* (Table 2). In addition, two compounds showed species-specific activity— one effective against *S. mansoni*, and the other against *S. japonicum* (Table 2). To assess their relative anti-schistosomal potency, IC50 values were calculated based on gradient concentrations and corresponding phenotype scores (Table 2, S4 and S5 Figs). Among these, HY-B2171A, predicted to target the splicing factor U2AF through virtual docking, exhibited the highest efficacy in both assays for *S. mansoni* and *S. japonicum*. Notably, IC50 analysis revealed significant variation in the compound’s efficacy between these two species. For instance, the compounds HY-B2171A, HY-15004, HY-100493, HY-15196, HY-18018, HY-14751, HY-12382 and HY-138407 exhibited greater efficacy against *S. mansoni* compared to *S. japonicum*. In contrast, compounds HY-112852, HY-12462, and HY-16322 demonstrated superior potency against *S. japonicum*. Notably, HY-100493 showed exclusive activity against *S. mansoni*, while HY-16322 displayed species-specific efficacy restricted to *S. japonicum*. In comparison to praziquantel, HY-B2171A exhibited stronger inhibitory effects on *S. mansoni* than on *S. japonicum*.

**Table 2.**
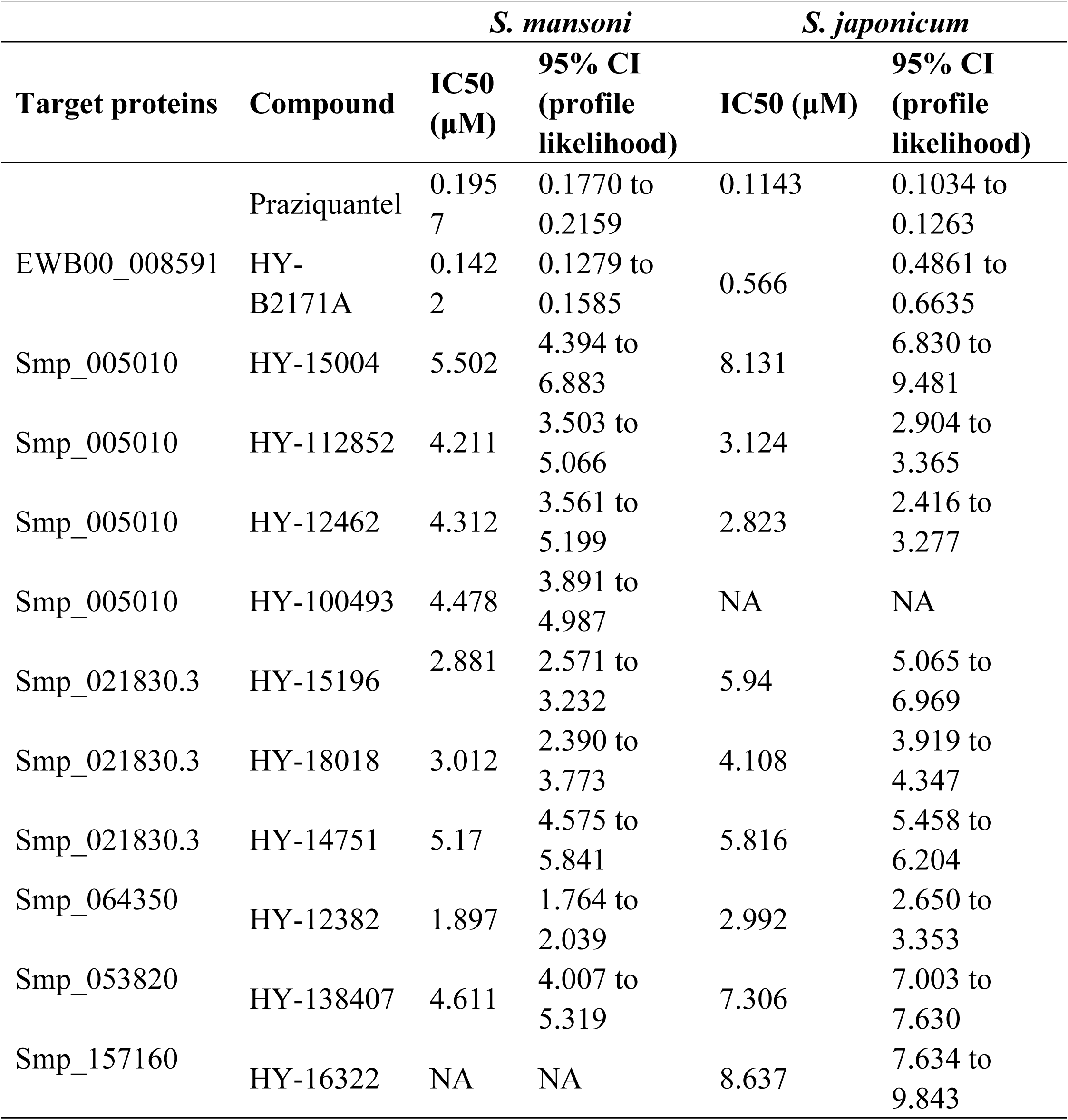
IC50 values for high-efficiency compounds tested in *S. mansoni* and *S. japonicum*.

### Characterization of the coding gene for CH’s potential target

CH was predicted to target the protein encoded by *EWB00_008591 in S. japonicum*, annotated as the splicing factor U2AF subunit (U2 small nuclear RNA auxiliary factor) (Fig 6E). Notably, this protein shares high sequence and structural similarity with Smp_019690 in *S. mansoni*, which is also annotated as a U2AF subunit (S5 Table, S2 Fig). Interestingly, CH also exhibited potent activity against *S. mansoni* (Tables 1 and 2). Given the high degree of similarity between the *Sj*U2AF and *Sm*U2AF, we further performed molecular docking of CH with Smp_019690, which resulted in similarly high docking scores (Boxed in Fig 6A). These findings suggest that CH may exert its anti-schistosome activity by targeting the conserved U2AF subunit in both species. Based on this hypothesis, we selected Smp_019690 as a candidate target for further functional and mechanistic studies in *S. mansoni*.

U2AF is a critical auxiliary component in RNA splicing, functioning as a heterodimer consisting of two subunits: U2AF65 and U2AF35 [21]. To classify the *S. mansoni* splicing factor U2AF subunit, we conducted a phylogenetic analysis, identifying it as part of the U2AF65 family (S6 Fig). Protein sequence alignments with homologs from other schistosome species and model organisms, including humans, mice, *Danio rerio*, and *Caenorhabditis elegans*, revealed conserved regions across these species (S7 Fig). These findings highlight evolutionary similarities and suggest the presence of functionally conserved domains in U2AF65.

As indicated by the single-cell atlas of *S. mansoni* (https://www.collinslab.org/schistocyte/), *Smp_019690* is predicted to be expressed across all cell types in both sexes (S8A and 8B Figs). Whole-mount *in situ* hybridization and fluorescent *in situ* hybridization further confirmed this, revealing higher expression levels in the sexual organs of the parasites, such as the testes and ovaries (S8C and 8D Figs).

### CH disrupts exon splicing in schistosomes by targeting U2AF65

To test whether CH inhibits schistosome activity by targeting *Sm*U2AF65 as predicted (Fig 6A), we first evaluated the performance of parasites subjected to *Smp_019690* RNAi or CH treatment. As expected, silencing the expression of *Smp_019690* via *in vitro* cultivation with dsRNA resulted in all worms detaching from the plate bottom and exhibiting significantly reduced motility by day 10 (Fig 7A). qPCR analysis showed that the expression level of *Smp_019690* was reduced by 66% (Fig 7B). When worms were treated with a low concentration (0.618 μM) of CH, all exhibited signs of morbidity within one day (Fig 7C). Given that U2AF65 has been previously reported to play a role in cell proliferation [22, 23], we further assessed the effect of RNAi or CH treatment on parasite cell proliferation using EdU labeling. Both treatments resulted in a significant decrease in proliferative cells in the soma and testes of male parasites (Figs 7A and 7C). To better understand the molecular mechanisms underlying these phenotypic changes. RNA-seq was performed to profile transcriptomic changes in both *Smp_019690*-knockdown adult male worms and CH-treated adult male worms (Fig 7D). Dramatic transcriptional changes occurred after both treatments (S9 Fig). Using selection criteria of *Padj* < 0.05 and |Log_2_FoldChange| ≥ 1, we identified 556 downregulated and 414 upregulated genes in the *Smp_019690* RNAi group compared to the control group (Fig 7E, S12 Table). In the CH-treated group compared to the DMSO group, 268 downregulated and 223 upregulated genes were identified (Fig 7F, S13 Table). A comparison of differentially expressed genes (DEGs) between the two groups revealed 145 shared genes (Fig 7G, S14 Table).

**Fig 7.**
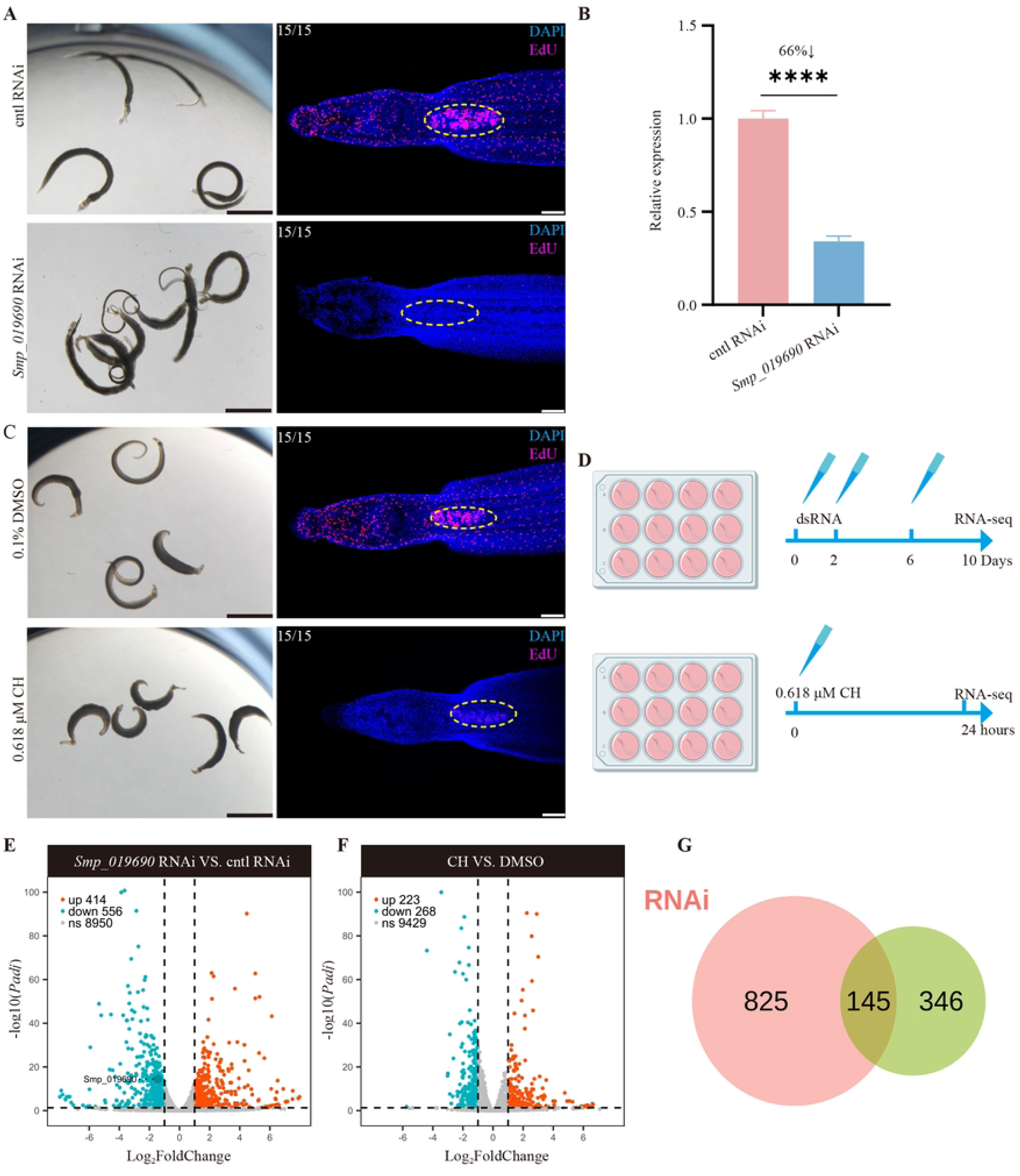
Stem cell maintenance and transcriptomic changes in parasites following *Smp_019690* knockdown or CH treatment. (A) Phenotype of adult *S. mansoni* following *Smp_019690* or control RNAi (left, n=15-18), and EdU labeling of proliferative cells in male parasites (right). Representative bright-field images from 3 biological replicates. Scale bars, 1000 μm. For EdU labelling, nuclei were stained with DAPI (blue) and proliferative cells were are shown in magenta. Testes are outlined with dashed yellow circles. n=15 worms from 3 biological replicates, Scale bars, 100 μm. (B) Relative mRNA expression levels of *Smp_019690* in parasites treated with *Smp_019690* dsRNA or control dsRNA (mean ± SD). *\*\*\*\*P*<0.0001. (C) Phenotype of adult *S. mansoni* following 0.618 μM CH or DMSO treatment (left), and EdU labeling of proliferative cells in male parasites (right). Representative bright-field images from 3 biological replicates. Scale bars, 1000 μm. For EdU labelling, nuclei were stained with DAPI (blue) and proliferative cells were are shown in magenta. Testes are outlined with dashed yellow circles. n=15 worms from 3 biological replicates, Scale bars, 100 μm. (D) Schematic diagram of sample collection for RNA-seq following RNAi and CH treatment. (E–F) Volcano plots of differentially expressed genes following Smp_019690 RNAi (E) or CH treatment (F). Differentially expressed genes were defined as those with *Padj* < 0.05 and |log₂FoldChange| ≥ 1. (g) Venn diagram of the differentially expressed genes between RNAi and CH treatment group.

In humans and plants, U2AF plays a key role in mediating alternative splicing [24-26]. A schematic representation of the working model for U2AF, based on these studies, is shown in Fig 8A. Specifically, U2AF35 recognizes and binds to the AG dinucleotide at the 3’ splice site, while U2AF65, which contains two RNA recognition motifs (RRM1/RRM2), interacts with the poly-pyrimidine tract (PPT) at the 3’ splice site [25]. Additionally, the C-terminal UHM (U2AF homology motif) of U2AF65 associates with the ULM (U2AF homology motif-like) domain of the auxiliary protein SF1 [24], stabilizing the early spliceosome complex. Together, U2AF35 and U2AF65 facilitate early spliceosome assembly, regulate exon inclusion or skipping, and influence the dynamic selection of splice sites [26]. Given the high sequence similarity between *Sm*U2AF65 and its human homologs, we hypothesized that *Sm*U2AF65 may also be involved in mRNA splicing in *S. mansoni*. To test this, we analyzed alternative splicing events using RNA-seq data from *Smp_019690* RNAi and CH treatment experiments (Figs 8B and 8C). Five types of alternative splicing events were assessed: skipped exon (SE), mutually exclusive exons (MXE), alternative 3’ splice sites (A3SS), retained intron (RI), and alternative 5’ splice sites (A5SS). Knocking down *Smp_019690* resulted in changes across all splicing event types, with SE events showing the greatest number of changes: 2,419 reduced and 201 increased events (Fig 8B). These findings suggest that *Sm*U2AF65 promotes SE events in *S. mansoni*, consistent with U2AF65’s role in other organisms (Fig 8A) [27]. Similarly, changes in splicing events were observed after treating worms with CH, a predicted binding compound for *Sm*U2AF65 (Fig 8C). Although the total number of altered splicing events was lower than in the RNAi group, SE events predominated. Notably, CH treatment resulted in an equal number of reduced and increased SE events, contrasting with the RNAi results where SE events mainly decreased.

**Figure 8.**
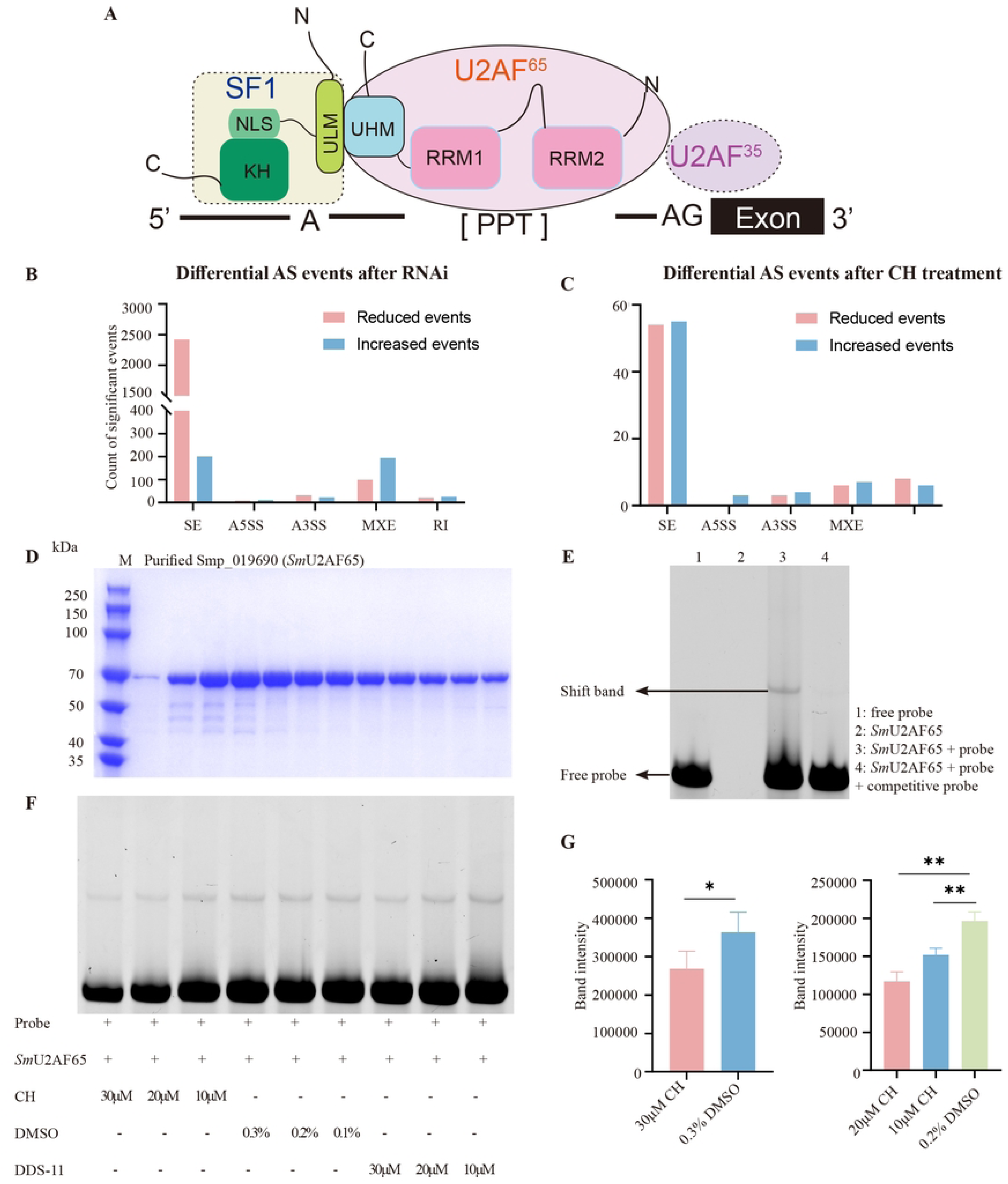
CH inhibits formation of schistosome U2AF65/RNA complex. (A) The model of U2AF-dependent splicing in model organisms [25]. NLS, nuclear localization signal; KH, K Homology Domain; ULM, U2AF Homology Motif-like; UHM, U2AF Homology Motif; RRM, RNA Recognition Motifs; PPT, the poly-pyrimidine tract. (B-C) Differential alternative splicing events (b) after RNAi or CH treatment (C). AS, alternative splicing; SE, skipped exon; A5SS, alternative 5’ splice sites; A3SS, alternative 3’ splice sites; MXE, mutually exclusive exons; RI, retained intron. (D) SDS-PAGE analysis of purified recombinant Smp_019690 (*Sm*U2AF65). Lane M: protein molecular weight marker; remaining lanes: purified Smp_019690 protein. (E) Assessment of the binding affinity between *Sm*U2AF65 and FAM-labeled RNA. (F) Assessment of the effect of varying CH concentrations on the binding interaction between *Sm*U2AF65 and RNA. DDS-11, an unrelated compound as control. (G) Quantitative analysis of *Sm*U2AF65/RNA shift band intensity at different CH concentrations. The quantitative analysis data are derived from Figure S10. The shift band was analyzed for grayscale intensity using the ImageJ software. *\*P*<0.05, *\*\*P*<0.01.

To explore the function of *Sm*U2AF65 in the parasite and its interaction with compound CH, we cloned and expressed the full-length *Sm*U2AF65 protein in *E. coli* (Fig 8D). The purified protein was then used for further analysis. As described previously [21, 26], U2AF65 plays a critical role in pre-mRNA splicing by recognizing and binding to the polypyrimidine tract near the 3’ splice site of introns, facilitate the assembly of the early spliceosome, regulate exon inclusion or skipping. Using rMATS, we identified the introns of the top 10 differentially skipped exons from our RNA-seq data of CH treatment experiments. After extracting and aligning these sequences with MAFFT, we identified a consensus 50 bp sequence (5’-UACAUAAAUUAAUAAGUAUUUUUACUUUUUUUUAUUUUCAAUUUCAACAG-3’) in the latter region, assumed to be the RNA fragment recognized by *Sm*U2AF65. This 50 bp RNA fragment was shown to bind to the purified *Sm*U2AF65, forming a shift band on native polyacrylamide gel, as detected by EMSA (Fig 8E). To determine whether CH disrupts the binding of U2AF65 to its target RNA, we conducted binding assays with *Sm*U2AF65, the RNA fragment, and varying concentrations of CH. The shift bands weakened as the CH concentration increased (Figs 8F and 8G, S10 Fig), whereas the unrelated compound DDS-11 had no effect. These findings suggest that CH targets U2AF65, disrupting the U2AF65/RNA complex and affecting RNA splicing in schistosomes.

In conclusion, we established a workflow combining highly accurate protein structure modeling and virtual docking to identify compounds binding to target proteins in schistosomes. Through *in vitro* screening, we prioritized compounds with high efficacy against adult and juvenile worms of *S. mansoni* and *S. japonicum*. We demonstrated that U2AF65 is a functional target, with CH identified as the most potent compound. CH treatment or U2AF65 knockdown resulted in significant changes in exon-skipping events, with CH inhibiting the binding of U2AF65 to its target RNA. This study provides a robust workflow for high-throughput screening of anti-schistosomal compounds and highlights a novel protein-compound interaction, accelerating the development of anti-schistosomal drugs.

## Discussion

Research on anti-schistosomal drugs dates back to 1918 with the discovery of potassium antimonyl tartrate [28]. Since then, continuous efforts have been made to develop effective treatments. A major breakthrough in schistosomiasis therapy was achieved in the mid-to-late 1970s with the discovery of praziquantel [29]. Despite ongoing research to develop new therapeutics, praziquantel remains the sole drug available for the treatment of this disease. In this study, we selected 202 target proteins essential for the viability of *S. mansoni* and *S. japonicum* from our previous RNAi screening. Among these, we prioritized 37 proteins with highly accurate predicted structures for virtual docking, leading to the identification of 268 compounds with high binding potential. Through *in vitro* cultivation and evaluation of parasite activity, we further identified 7 distinct compounds that exhibit significant inhibitory effects at 1 μM on *S. mansoni* and *S. japonicum*. Among them, HY-B2171A, which targets schistosome U2AF65, demonstrated the most robust efficacy against both species. This workflow provides a promising approach to accelerate the development of alternative drugs to praziquantel.

During the *in vitro* evaluation of the anti-parasite efficacy, we observed distinct susceptibilities between adult and juvenile worms to the same compound. Specifically, compounds HY-12844, HY-112647 and HY-N0794 demonstrated higher efficacy against *S. mansoni* adults than juveniles, whereas compounds HY-138696, HY-15002, HY-B0119 and HY-11101 were more effective against juveniles than adults (S6 Table). A similar pattern was observed in *S. japonicum*. This differential susceptibility between developmental stages was further exemplified by the anti-schistosomal effects of PQZ, where stage-specific efficacy variations were particularly evident. Previous studies have reported that juvenile *S. mansoni* (3–4 weeks post-infection) and *S. japonicum* (2 weeks post-infection) are not susceptible to PZQ *in vivo* [30-32], preventing the treatment of recent infections [33]. However, the mechanisms underlying this reduced susceptibility remain unclear. Investigations into oxamniquine derivatives further highlighted that two novel compounds exhibited weaker activity against juveniles compared to adults [34]. Similarly, compounds such as artemether and curcumin have demonstrated preferential efficacy against juveniles [35]. Collectively, these findings indicate that differential compound susceptibility arises from physiological and structural divergences between juvenile and adult worms. This stage-specific susceptibility is not restricted to individual compounds but represents a critical factor that must be systematically considered in anti-schistosomal drug discovery.

Furthermore, our work revealed marked interspecies variations in compound efficacy against different *Schistosoma* species. Compounds HY-B2171A, HY-15004, HY-100493, HY-15196, HY-18018, HY-14751, HY-12382 and HY-138407 exhibited significantly lower IC50 values against *S. mansoni* compared to *S. japonicum*. Conversely, HY-112852, HY-12462, HY-16322 demonstrated superior potency against *S. japonicum* (Table 2). Notably, HY-100493 displayed exclusive activity against *S. mansoni*, while HY-16322 showed species-specific efficacy restricted to *S. japonicum*. This pharmacological divergence aligns with previous findings. A comprehensive review by Abdul-Ghani et al. highlighted oxamniquine’s exclusive activity against *S. mansoni* [36]. Similarly, prior studies have demonstrated that CDDD-0149830 exhibits greater efficacy against *S. haematobium*, CIDD-150303 shows superior activity against *S. mansoni*, and CIDD-066790 is more effective against *S. japonicum* [34]. The observed differences in compound efficacy across *Schistosoma* species may stem from variations in parasite biology. Further investigation is warranted to elucidate the precise mechanisms underlying these interspecies pharmacological discrepancies.

Structure-based drug discovery plays a crucial role in identifying anti-schistosomal compounds by leveraging the 3D structures of target proteins to facilitate precise compound screening, improve hit rates, and reduce the workload and cost of experimental screening. In this study, we employed AlphaFold to predict the structures of target proteins, followed by molecular docking to assess interactions between the target proteins and small-molecule compounds, aiming to identify potential anti-schistosomal candidates. A key strength of this approach is that AlphaFold generates high-confidence 3D structure predictions for proteins lacking experimentally resolved structures, thereby expanding the range of schistosomal targets available for structure-based drug screening. However, this method has inherent limitations. First, AlphaFold predictions rely on sequence homology and physicochemical properties, which may not fully capture dynamic conformational changes, particularly the plasticity of ligand-binding pockets, potentially affecting virtual screening accuracy. Second, if a target protein is highly conserved between the host and schistosomes, the selected compounds may carry a risk of host toxicity. To mitigate this, structural comparisons should be performed to identify parasite-specific binding sites for further optimization. For schistosome-specific or poorly conserved targets, screened compounds are more likely to selectively inhibit the parasites without interfering with host physiological functions, thereby reducing toxicity risks. However, many schistosome-specific proteins have poorly characterized functions, and the absence of experimentally determined structures and ligand-binding data lowers confidence in AlphaFold-predicted models, necessitating experimental validation. Nonetheless, protein expression and structural determination remain significant challenges. Thus, in practical applications, SBDD based on AlphaFold-predicted structures should be integrated with target conservation analysis, molecular dynamics simulations, and experimental validation to enhance screening efficiency and improve the success rate of identifying selective anti-schistosomal drugs.

In this study, we found compound CH displayed a strongly inhibitory effect on schistosome viability *in vitro*. Virtual docking analysis predicted its potential target as U2AF65, a key protein involved in pre-mRNA splicing [26]. Transcriptomic analysis revealed significant differences between samples treated with CH and those subjected to RNAi of the target gene (Figs 7E and F). We hypothesize that this discrepancy arises from the fundamental differences in their mechanisms of action: CH directly targets the proteins, whereas RNAi affects gene expression. Specifically, RNAi reduces U2AF65 protein levels by degrading its mRNA, gradually impairing RNA splicing. In contrast, CH directly targets the U2AF65 protein to disrupt its function. Alternative splicing analysis further confirmed that CH affects splicing events in a manner similar to, but not entirely identical to, U2AF65 knockdown, as both treatments significantly influenced exon skipping (Figs 8B and C). Previous studies have shown that U2AF65 plays a crucial role in pre-mRNA splicing by recognizing and binding to the polypyrimidine tract near the 3’ splice site of introns, facilitating early spliceosome assembly and regulating exon inclusion or skipping [21, 26]. Our findings support this established mechanism, as *in vitro* evaluation confirmed that U2AF65 binds to a polypyrimidine RNA probe.

CH, a member of the Bioactive Compound Library, is a microbially derived compound previously reported to target the Golgi complex, inducing apoptosis in clear cell renal carcinoma cells [37, 38]. It also inhibits the template activity of DNA in the DNA-dependent RNA polymerase system by complexing with DNA [39] and acts as a potent inhibitor of VHL-deficient (VHL⁻/⁻) CCRCC cell proliferation [38]. In this study, we found that both CH treatment and *U2AF65* knockdown significantly suppressed parasite cell proliferation (Figs 7A and 7C), consistent with previous findings [22, 23, 38]. Our results demonstrate that CH interferes with RNA splicing in schistosomes by inhibiting U2AF65 ‘s RNA-binding ability, leading to stem cell death and ultimately resulting in parasite death. While U2AF65 inhibition alone is sufficient to induce parasite lethality, CH may also interact with additional targets in schistosomes. Despite its promising anti-schistosomal activity, the high sequence similarity between schistosome U2AF65 and its human homolog raises concerns about potential host toxicity. Therefore, rational structural modifications of CH are recommended to enhance its binding selectivity toward the parasite protein over the human counterpart, thereby improving its safety and therapeutic index.

Moreover, the integrated strategy employed in this study—combining AlphaFold-based structural prediction, virtual screening, and functional validation—holds significant promise for application to other parasitic diseases, such as malaria or leishmaniasis, where high-resolution structural data for protein targets remain scarce. This approach may accelerate target prioritization and compound discovery across a broader range of neglected tropical diseases.

## Materials and Methods

### Ethics statement

All Experiments involving vertebrate animals were conducted in compliance with protocols approved by the Institutional Animal Care and Use Committee (IACUC) of Fudan University (approval APN: 2021JS0078).

### Parasites

Adult *S. mansoni* (6–7 weeks post-infection) and juvenile stage worms (4 weeks post-infection) were harvested from infected female Kunming (KM) mice by hepatic portal vein perfusion using ice-chilled 0.9% (w/v) NaCl containing 0.005% (w/v) heparin. For *S. japonicum*, adult worms (4 weeks post-infection) and juvenile stage worms (2 weeks post-infection) were used. Parasites were rinsed in DMEM containing 10% Horse Serum, then cultured in DMEM (high glucose) supplemented with 10% fetal bovine serum and 1× Antibiotic-Antimycotic. Juvenile worms were cultured in Basch’s medium 169 [40]. The host mice utilized in the experiment were 4-week-old female Kunming mice (KM), weighing between 18 and 22 g. The mice were maintained under standard laboratory conditions, with the temperature controlled at 22 ± 2°C, relative humidity maintained at 50-60%, and a 12-hour light/dark cycle.

### Virtual screening workflow for protein-ligand docking

The 3D structures of proteins were retrieved from the AlphaFold database (https://www.alphafold.ebi.ac.uk/). After filtering out low-quality models with an average per-residue model confidence score (pLDDT) < 70, high-quality protein models were retained for further analysis. Using Schrödinger Maestro 11.4, hydrogen atoms were added to the protein models, followed by energy optimization using the OPLS2005 force field (RMSD = 0.30 Å). The binding sites of the protein models were predicted using the Sitemap tool, and the best site (Site1) was selected as the center of a docking box (20 Å × 20 Å × 20 Å).

To identify potential active compounds targeting these proteins, the MCE Bioactive Compound Library Plus (14,600 compounds) was used as the source for candidate compounds. The 2D structures of the compounds were processed by adding hydrogen atoms and performing energy optimization, following the same procedure as for the protein models, to generate 3D structures for docking.

Ligand docking was performed using the compound library against each protein model in three stages: first, High-Throughput Virtual Screening (HTVS) was conducted to identify primary compound candidates; next, the top 10% of compounds from the HTVS results, based on docking scores, were subjected to Standard Precision (SP) docking; finally, the top 10% of compounds from the SP results underwent a final round of docking using Extra Precision (XP) mode. After these steps, the top 200 compounds with the highest docking scores were selected as candidates for further analysis.

### Sequence and structural comparison of target proteins

The protein sequence of *S. mansoni* was obtained from the *Sm*.V7 genome assembly, while the sequence of *S. japonicum* was retrieved from the PRJNA520774 - HuSjv2 dataset. Sequence alignment of target proteins from *S. mansoni* and *S. japonicum* was conducted using the WormBase Parasite database to determine sequence similarity. For structural comparison, predicted protein structures were sourced from the AlphaFold Protein Structure Database. Structural alignments of homologous proteins from *S. mansoni* and *S. japonicum* were performed using PyMOL, and root mean square deviation (RMSD) values were calculated to quantitatively assess structural similarity.

### Evaluation of compound effects on parasites

*In vitro* evaluation of the selected compounds (at a single concentration of 10 μM in 0.1% DMSO) was conducted on adult worms (4–6 pairs in 12-well plates with 3 mL medium, or 2 pairs in 24-well plates with 1 mL medium) and juvenile worms (10–16 individuals in 48-well plates with 500 μL medium). Worms were cultured for 7 days at 37°C in a humidified atmosphere with 5% CO₂, with the medium replaced every 2 days. Phenotypes were recorded daily under a microscope. Control groups included worms treated with 0.1% DMSO (negative control) and 1 μM PZQ (positive control) in DMSO. At day 7, microscopy images were captured using a digital camera system (OLYMPUS, SZX2-ILLTS).

To quantify adult worm attachment to the substrate following compound treatment, freshly perfused adult worms were placed in 24-well plates with 1 mL media and cultured overnight. On the following day (D1), unattached worms were removed, and compounds were added to the media. The media and compounds were replaced on D1, D3, and D5. Dose-response titration (25 μM–0.010 μM) was performed for effective compounds to assess their anti-schistosomal potency. Eight titration points with two-fold or three-fold stepwise dilutions from the top concentration were prepared, and each titration point was performed in triplicate (4 male worms per replicate). Short videos (5 seconds) were recorded under a microscope at day 7 to facilitate scoring. Compound effects were evaluated using 3 indicators [41]: 1) attachment: 1 = attached, 0 = unattached; 2) motility: 0 = static, 1 = slight drifting, 2 = intermittent swinging, 3 = energetic swinging and 3) morphological changes: 0 = tegumental disruption or dark worms, 1 = normal appearance). Worm viability was scored based on the sum of these three indicators, with a scale ranging from 0 to 5. Data on phenotypes and viability scores from the dose-response titration were analyzed using GraphPad Prism to generate approximate IC50 values for each effective compound.

### Parasite staining and imaging

Whole-mount *in situ* hybridization (WISH), fluorescent *in situ* hybridization (FISH) [42], DAPI staining, and EdU labeling [43] were performed as previously described. Worms stained with WISH were imaged using light microscopy (Axio Zoom.V16, ZEISS), while fluorescent images were captured with confocal scanning microscopy (A1, Nikon). Riboprobe synthesis for WISH/FISH: Primers were designed using the Primer-BLAST tool (https://www.ncbi.nlm.nih.gov/tools/primer-blast/index.cgi?LINK_LOC=BlastHome) based on the coding sequence (CDS) of the target gene to amplify fragments ranging from 400 to 700 bp, as listed in S1 Table. Sequence specificity was carefully analyzed to ensure accuracy. The PCR products were inserted into the pJC53.2 vector using TA cloning. The resulting plasmids were purified from *E. coli*, validated by sequencing, and then used as templates for riboprobe generation.

### dsRNA synthesis

dsRNA production was conducted as previously described [44, 45]. Briefly, the recombinant pJC53.2 plasmid, containing the target gene fragment as previously described, was utilized as a template for dsRNA synthesis. The target dsRNA was synthesized *in vitro* using a T7 High Yield RNA Synthesis Kit (Shanghai Yeasen BioTechnologies, Chian) for RNAi applications. The oligonucleotide sequences used for dsRNA template generation are provided in S1 Table.

### Sample collection for RNA-seq

To assess gene expression changes following the knockdown of *Smp_019690*, 6 adult male worms were placed in each well of a 12-well plate with 3 mL of Basch 169 medium, supplemented with 30 µg/mL dsRNA. The medium was replaced every two days, and dsRNA was administered at day 0, 2, and 6. Worms were collected at day 10 for sequencing. As controls, worms cultured in parallel were treated with dsRNA derived from a bacterial gene [44]. To evaluate gene expression changes following treatment with the compound CH, 6 adult male worms were placed in each well of a 12-well plate with 3 mL of Basch 169 medium. Parasites were treated with 0.618 μM CH for 24 hours, with worms treated with DMSO as control. After *Smp_019690* RNAi or CH treatment, worms were harvested and washed with PBS, and then flash frozen in liquid nitrogen for RNA extraction and Illumina sequencing (Novogene, China). For both experiments, each biological replicate consisted of worms from two wells, with three biological replicates for each condition.

### RNA-seq data processing and splicing analysis

Schistosome total RNA was isolated using Qiagen miRNeasy Mini Kit and qualified using an Agilent 5400 system. RNA sequencing was performed according to the manufacturer’s protocol on Illumina platforms with PE150 strategy in Novogene Bioinformatics Technology Co., Ltd (Beijing, China). For differential expression analysis, raw sequencing reads were aligned to the reference genome using Hisat2 [46], a fast and sensitive splice-aware aligner optimized for RNA-seq data. The resulting alignment files were subsequently processed with StringTie [47] to assemble transcripts and quantify gene-level expression. The generated count matrix was then used as input for DESeq2 [48], a widely utilized R package that employs a negative binomial distribution model to identify differentially expressed genes (DEGs). DESeq2 performed data normalization, dispersion estimation, and statistical testing to detect significant expression changes between experimental conditions. Genes with an adjusted *P*-value < 0.05 and |Log_2_FoldChange| ≥ 1 were considered significantly differentially expressed.

For alternative splicing analysis, we employed rMATS (replicate Multivariate Analysis of Transcript Splicing) [49], a robust computational tool for detecting differential alternative splicing events from RNA-seq data. BAM files generated by StringTie were used as input for rMATS, and the analysis was conducted with default parameters. This approach enabled the identification of five major types of alternative splicing events: skipped exons (SE), retained introns (RI), mutually exclusive exons (MXE), and alternative 5’ and 3’ splice sites (A5SS and A3SS). rMATS implemented a statistical framework to compare splicing patterns between experimental groups and estimated the inclusion level (ψ values) for each splicing event. Significance was assessed using false discovery rate (FDR)-corrected *P*-values with threshold of 0.05.

### Expression and purification of recombinant Smp_019690

The splicing factor U2AF subunit (Smp_019690) of *S. mansoni* was expressed in *E.coli* strain ArcticExpress (DE3) as a His+SUMO fusion protein in the pET28a vector. The recombinant protein was purified using Ni^2+^ affinity chromatography on a HisTrap column (Cytiva), followed by size-exclusion chromatography on a Superdex-75 prep-grade column (GE Healthcare). The oligonucleotide sequences used to generate the full-length *Smp_019690* template were provided in S1 Table.

The recombinant protein was initially bound to a HisTrap column equilibrated with a buffer containing 20 mM Tris-HCl, 300 mM NaCl, and 20 mM imidazole at pH 8.0. Elution was performed using a buffer containing 20 mM Tris-HCl, 300 mM NaCl, and 500 mM imidazole at pH 8.0. Protein purification was carried out using the AKTA protein purification system, and elution was monitored via UV detection at 280 nm to track the protein in the eluted fractions. Purity was confirmed by SDS-PAGE gel analysis, which was stained with Coomassie Brilliant Blue.

To cleave the His+SUMO tag, the recombinant protein was treated with ULP1 protease during dialysis against a buffer containing 20 mM Tris·HCl, 300 mM NaCl, pH 8.0, 10% (v/v) glycerol, and 0.1 mM EDTA. The cleaved His+SUMO tag was separated from U2AF65 using Ni²⁺ affinity chromatography, followed by size-exclusion chromatography on a previously equilibrated Superdex-75 prep-grade column with 20 mM Tris·HCl, 300 mM NaCl, pH 8.0, and 0.1 mM EDTA. The purified U2AF65 was concentrated using an Ultra Centrifugal Filter (Millipore) with a 50 kDa MWCO, and protein concentration was estimated by its extinction coefficient (38,655 M⁻¹cm⁻¹) and absorbance at 280 nm.

### Electrophoretic mobility shift assay (EMSA) for protein/RNA binding

FAM labeled and unlabeled RNA fragment were synthesized (Genscript, China) and diluted to a final concentration of 100 μM in TE buffer (10 mM Tris-HCl, 0.1 mM EDTA, pH 8.0). The RNA fragments were denatured by incubating at 70°C for 5 minutes, followed by immediate cooling on ice. Half pmol of FAM-labeled RNA was incubated with 1 μg of recombinant U2AF65 protein in reaction buffer (Beyotime, China) for 20 minutes at room temperature, in a final volume of 10 μL. For the competitive control group, 200 pmol of unlabeled RNA was added. In the compound-binding groups, the compounds (10-30 μM) were pre-incubated with the protein for 30 minutes at room temperature before adding the RNA fragment. The reactions were analyzed by electrophoresis on 6% non-denaturing polyacrylamide gels, prepared in 1x TBE buffer (89 mM Tris, 89 mM boric acid, 2 mM EDTA, pH 8.3). A pre-run was performed at 200 V for 10 minutes, followed by electrophoresis at 200 V for approximately 25 minutes. The gel was visualized using a FLA Typhoon 9000 (GE), and signals from the entire lanes were quantified using ImageJ software.

### Statistical analysis

Statistical analysis of data was performed using GraphPad Prism software, and results are presented as mean ± SD. Statistical significance was determined using an unpaired two-tailed parametric *t*-test, where *P* values < 0.05 were considered significant. The significance levels are indicated as follows: NS (not significant); *\*P* < 0.05; *\*\*P* < 0.01; *\*\*\*P* < 0.001; *\*\*\*\*P* < 0.0001.

## Data sharing statement

The raw sequencing data reported in this paper have been deposited in the NCBI Sequence Read Archive (SRA) database under the accession number PRJNA1264060.

## Acknowledgments

We thank Professor Jinbiao Ma and his team at the Department of Biochemistry and Biophysics, School of Life Sciences, Fudan University, for their valuable assistance with protein purification.

This study was supported by grants from the National Key Research and Development Program of China (Grant No. 2021YFC2300800), the Fund of Fudan University and Cao’ejiang Basic Research (No. 24FCA04), and the Science and Technology Leading Talent Team in Inner Mongolia Autonomous Region (No. 2022LJRC0009). Animations created with BioGDP.com.

## Supporting information

**S1 Fig. RNAi-induced phenotypes of target genes in *S. mansoni* and *S. japonicum*.**

(A) Bright-field images showing phenotypes of adult *S. mansoni* after RNAi treatment. (B) Bright-field images showing representative phenotypes of adult *S. japonicum* after RNAi treatment. Representative images from 3 biological replicates, Scale bars, 1000 μm.

**S2 Fig. Structural alignment of homologs between *S. mansoni* and *S. japonicum* for 37 target proteins.**

Each alignment demonstrates the structural similarity between homologous proteins from the two schistosome species. Protein structures were superimposed using PyMOL, and RMSD values were calculated to quantify structural similarity. The protein structure of *S. mansoni* is shown in cyan, while that of *S. japonicum* is depicted in salmon. The small inset in the lower left corner represents the alignment of the full-length protein structures, whereas the enlarged section highlights the alignment of conserved regions or specific protein domains.

**S3 Fig. Phenotypic details of adult worms treated with lethal compounds.**

(A) Phenotypes of *S. mansoni* treated with different compounds at concentration of 10 μM for 7 days. (B) Phenotypes of *S. japonicum* treated with different compounds at concentration of 10 μM for 7 days. DMSO as negative control; PZQ as positive control. Scale bars, 500 μm. Red arrows denote tegumental erosion.

**S4 Fig. Dose–responses analysis of highly effective compounds in *S. mansoni*.** IC50 values were determined for ten high-efficacy compounds targeting adult *S. mansoni*. The fitted IC50 curves are shown in the figure, with praziquantel (PZQ) used as a control. Dose-response curves were generated using GraphPad Prism based on IC50 values calculated from experimental data. The IC50 value was evaluated on Day 7 (D7) post-treatment.

**S5 Fig. Dose–responses analysis of highly effective compounds in *S. japonicum*.** IC50 values were determined for ten high-efficiency compounds targeting adult *S. japonicum.*, The fitted IC50 curves are shown in the figure, with praziquantel (PZQ) used as a control. Dose-response curves were generated using GraphPad Prism based on IC50 values calculated from experimental data. The IC50 value was evaluated on Day 7 (D7) post-treatment.

**S6 Fig. Phylogenetic analysis of the protein product of *Smp_019690*.**

The green leaf represents U2AF35, the red leaf represents U2AF65. The tree was constructed using the Maximum Likelihood method in MEGA 11, based on amino acid sequences from 7 species. Bootstrap values from 1,000 replicates are indicated at the nodes. Smp_019690 belongs to the U2AF65 family.

**S7 Fig. Multiple sequence alignment of U2AF65 protein.**

Multiple sequence alignment of U2AF65 proteins across various species, including *S. mansoni* (Smp_019690.1), *S. haematobium* (MS3_00003258.1), *S. japonicum* (EWB00_008591), *Caenorhabditis elegans* (NP_001022967.1), *Danio rerio* (NP_991252.1), *Mus musculus* (NP_001192160.1), and *Homo sapiens* (NP_009210.1). The percentage of conserved residues was calculated for each column based on physicochemical properties. Columns with a similarity score above 0.7 are considered highly conserved, with residues highlighted in red and framed in blue. Strictly identical residues appear in white on a red background. The blue-font-labeled line above the alignment represents the predicted secondary structure of human U2AF65, while the blue-font-labeled line below the alignment corresponds to that of *S. mansoni* U2AF65.

**S8 Fig. Localization of *Smp_019690* expression in *S. mansoni*.**

(A, B) UMAP plots showing the expression pattern of *Smp_019690* in adult female (A) and adult male (B) *S. mansoni*, obtained from the *S. mansoni* single-cell atlas (https://www.collinslab.org/schistocyte/). Color intensity represents the expression levels, with darker shades indicating higher expression. (C) WISH showing expression of *Smp_019690* in adult female and male worms. Representative images from n>5 worms. Scale bars, 100 μm. (D) FISH showing expression of *Smp_019690* in somatic tissues and reproductive organs of adult female and male worms. Representative images from n>5 worms. Nuclei are labelled by DAPI (grays). Scale bars, 50 μm.

**S9 Fig. Transcriptomic profiles of male schistosomes under *Smp_019690* RNAi and CH treatment.**

(A) PCA on the RNA-seq data under *Smp_019690* RNAi. (B) Heatmap of the Euclidean distance matrix with hierarchical clustering of transcriptomes from *Smp_019690* RNAi and control RNAi samples (C) Heatmap showing the cluster of top 2000 highly variable genes in the parasites from *Smp_019690* RNAi and control group. (D) PCA on the RNA-seq data under CH treatment. (E) Heatmap of the Euclidean distance matrix with hierarchical clustering of transcriptomes from CH and DMSO treatment samples. (F) Heatmap showing the cluster of top 2000 highly variable genes in the parasites from CH and DMSO treatment group.

**S10 Fig. EMSA analysis of *Sm*U2AF65 binding activity at varying CH concentrations.** (A, B) Effects of varying concentrations of CH on the binding interaction between *Sm*U2AF65 and RNA. Grayscale intensity analysis of the shift band was performed using ImageJ software for quantitative evaluation.

**S1 Table. Oligonucleotides used for RNAi, protein expression, *in situ* hybridization and qPCR.**

**S2 Table. Target proteins for virtual docking.**

**S3 Table. Potential binding compounds for each target protein from MCE library.**

**S4 Table. Selected 268 compounds against 37 target proteins.**

**S5 Table. Comparison of target protein sequences and structures between *S. mansoni* and *S. japonicum*.**

**S6 Table. Information of effective compounds against *S. mansoni* and *S. japonicum* at 10 μM.**

**S7 Table. Phenotypes of adult *S. mansoni* under the treatment of effective compounds**

**S8 Table. Phenotypes of effective compounds against *S. mansoni* and *S. japonicum* at 1 μM.**

**S9 Table. Phenotypes of adult *S. japonicum* under the treatment of effective compounds.**

**S10 Table. Phenotypes of juvenile *S. mansoni* under the treatment of effective compounds.**

**Table S11. Phenotypes of juvenile *S. japonicum* under the treatment of effective compounds.**

**Table S12. List of DEGs identified in the *Smp_019690* RNAi group vs. control group.**

**Table S13. List of DEGs identified in the CH-treated group vs. DMSO group.**

**Table S14. List of overlapping DEGs between RNAi and CH treatment groups**

## References

1. Lackey EK, Horrall S. Schistosomiasis. StatPearls. Treasure Island (FL) ineligible companies. Disclosure: Shawn Horrall declares no relevant financial relationships with ineligible companies.2024.

2. Tan SY, Ahana A. Theodor Bilharz (1825-1862): discoverer of schistosomiasis. Singapore Med J. 2007;48(3):184-5. PubMed PMID: 17342284.

3. WHO. Schistosomiasis. Available from: https://www.who.int/news-room/fact-sheets/detail/schistosomiasis.

4. Nelwan ML. Schistosomiasis: Life Cycle, Diagnosis, and Control. Curr Ther Res Clin Exp. 2019;91:5–9. doi: 10.1016/j.curtheres.2019.06.001. PubMed PMID: 31372189; PubMed Central PMCID: PMCPMC6658823.

5. King CH, Sutherland LJ, Bertsch D. Systematic Review and Meta-analysis of the Impact of Chemical-Based Mollusciciding for Control of *Schistosoma mansoni* and *S. haematobium* Transmission. PLoS Negl Trop Dis. 2015;9(12):e0004290. doi: 10.1371/journal.pntd.0004290. PubMed PMID: 26709922; PubMed Central PMCID: PMCPMC4692485.

6. Cioli D, Pica-Mattoccia L, Basso A, Guidi A. Schistosomiasis control: praziquantel forever? Mol Biochem Parasitol. 2014;195(1):23–9. doi: 10.1016/j.molbiopara.2014.06.002. PubMed PMID: 24955523.

7. Abdulla MH, Ruelas DS, Wolff B, Snedecor J, Lim KC, Xu F, et al. Drug discovery for schistosomiasis: hit and lead compounds identified in a library of known drugs by medium-throughput phenotypic screening. PLoS Negl Trop Dis. 2009;3(7):e478. doi: 10.1371/journal.pntd.0000478. PubMed PMID: 19597541; PubMed Central PMCID: PMCPMC2702839.

8. Moreira-Filho JT, Silva AC, Dantas RF, Gomes BF, Souza Neto LR, Brandao-Neto J, et al. Schistosomiasis Drug Discovery in the Era of Automation and Artificial Intelligence. Front Immunol. 2021;12:642383. doi: 10.3389/fimmu.2021.642383. PubMed PMID: 34135888; PubMed Central PMCID: PMCPMC8203334.

9. Kuntz AN, Davioud-Charvet E, Sayed AA, Califf LL, Dessolin J, Arner ES, et al. Thioredoxin glutathione reductase from *Schistosoma mansoni*: an essential parasite enzyme and a key drug target. PLoS Med. 2007;4(6):e206. doi: 10.1371/journal.pmed.0040206. PubMed PMID: 17579510; PubMed Central PMCID: PMCPMC1892040 manuscript and none of the authors have competing financial interests. There is no significant overlap between the submitted manuscript and any other papers from the same authors under consideration or in press elsewhere.

10. Long T, Neitz RJ, Beasley R, Kalyanaraman C, Suzuki BM, Jacobson MP, et al. Structure-Bioactivity Relationship for Benzimidazole Thiophene Inhibitors of Polo-Like Kinase 1 (PLK1), a Potential Drug Target in *Schistosoma mansoni*. PLoS Negl Trop Dis. 2016;10(1):e0004356. doi: 10.1371/journal.pntd.0004356. PubMed PMID: 26751972; PubMed Central PMCID: PMCPMC4709140.

11. Fioravanti R, Mautone N, Rovere A, Rotili D, Mai A. Targeting histone acetylation/deacetylation in parasites: an update (2017-2020). Curr Opin Chem Biol. 2020;57:65–74. doi: 10.1016/j.cbpa.2020.05.008. PubMed PMID: 32615359.

12. Kalinin DV, Jana SK, Pfafenrot M, Chakrabarti A, Melesina J, Shaik TB, et al. Structure-Based Design, Synthesis, and Biological Evaluation of Triazole-Based smHDAC8 Inhibitors. ChemMedChem. 2020;15(7):571–84. doi: 10.1002/cmdc.201900583. PubMed PMID: 31816172; PubMed Central PMCID: PMCPMC7187165.

13. Jilkova A, Horn M, Fanfrlik J, Kuppers J, Pachl P, Rezacova P, et al. Azanitrile Inhibitors of the SmCB1 Protease Target Are Lethal to *Schistosoma mansoni*: Structural and Mechanistic Insights into Chemotype Reactivity. ACS Infect Dis. 2021;7(1):189–201. doi: 10.1021/acsinfecdis.0c00644. PubMed PMID: 33301315; PubMed Central PMCID: PMCPMC7802074.

14. Marchant JS, Harding WW, Chan JD. Structure-activity profiling of alkaloid natural product pharmacophores against a Schistosoma serotonin receptor. Int J Parasitol Drugs Drug Resist. 2018;8(3):550–8. doi: 10.1016/j.ijpddr.2018.09.001. PubMed PMID: 30297303; PubMed Central PMCID: PMCPMC6287472.

15. Liu J, Dyer D, Wang J, Wang S, Du X, Xu B, et al. 3-oxoacyl-ACP reductase from *Schistosoma japonicum*: integrated in silico-in vitro strategy for discovering antischistosomal lead compounds. PLoS One. 2013;8(6):e64984. doi: 10.1371/journal.pone.0064984. PubMed PMID: 23762275; PubMed Central PMCID: PMCPMC3676400.

16. Wang J, Paz C, Padalino G, Coghlan A, Lu Z, Gradinaru I, et al. Large-scale RNAi screening uncovers therapeutic targets in the parasite *Schistosoma mansoni*. Science. 2020;369(6511):1649-53. doi: 10.1126/science.abb7699. PubMed PMID: 32973031; PubMed Central PMCID: PMCPMC7877197.

17. Wuthrich K. Protein structure determination in solution by NMR spectroscopy. J Biol Chem. 1990;265(36):22059–62. PubMed PMID: 2266107.

18. Shi Y. A glimpse of structural biology through X-ray crystallography. Cell. 2014;159(5):995–1014. doi: 10.1016/j.cell.2014.10.051. PubMed PMID: 25416941.

19. Earl LA, Falconieri V, Milne JL, Subramaniam S. Cryo-EM: beyond the microscope. Curr Opin Struct Biol. 2017;46:71–8. doi: 10.1016/j.sbi.2017.06.002. PubMed PMID: 28646653; PubMed Central PMCID: PMCPMC5683925.

20. Yang Z, Zeng X, Zhao Y, Chen R. AlphaFold2 and its applications in the fields of biology and medicine. Signal Transduct Target Ther. 2023;8(1):115. doi: 10.1038/s41392-023-01381-z. PubMed PMID: 36918529; PubMed Central PMCID: PMCPMC10011802.

21. Glasser E, Maji D, Biancon G, Puthenpeedikakkal AMK, Cavender CE, Tebaldi T, et al. Pre-mRNA splicing factor U2AF2 recognizes distinct conformations of nucleotide variants at the center of the pre-mRNA splice site signal. Nucleic Acids Res. 2022;50(9):5299–312. doi: 10.1093/nar/gkac287. PubMed PMID: 35524551; PubMed Central PMCID: PMCPMC9128377.

22. Yu Y, Zhen Z, Qi H, Yuan X, Gao X, Zhang M. U2AF65 enhances milk synthesis and growth of bovine mammary epithelial cells by positively regulating the mTOR-SREBP-1c signalling pathway. Cell Biochem Funct. 2019;37(2):93–101. doi: 10.1002/cbf.3378. PubMed PMID: 30773658.

23. Zhu Y, Song D, Guo J, Jin J, Tao Y, Zhang Z, et al. U2AF1 mutation promotes tumorigenicity through facilitating autophagy flux mediated by FOXO3a activation in myelodysplastic syndromes. Cell Death Dis. 2021;12(7):655. doi: 10.1038/s41419-021-03573-3. PubMed PMID: 34183647; PubMed Central PMCID: PMCPMC8238956.

24. Wang EJ, Kim YC, Lee JH, Kim JK. Identification of a nuclear localization signal mediating the nuclear import of splicing factor1. Plant Biotechnol Rep. 2021;15(6):775–81. doi: 10.1007/s11816-021-00722-0. PubMed PMID: WOS:000716854500001.

25. Fukumura K, Yoshimoto R, Sperotto L, Kang HS, Hirose T, Inoue K, et al. SPF45/RBM17-dependent, but not U2AF-dependent, splicing in a distinct subset of human short introns. Nat Commun. 2021;12(1):4910. doi: 10.1038/s41467-021-24879-y. PubMed PMID: 34389706; PubMed Central PMCID: PMCPMC8363638.

26. Shao C, Yang B, Wu T, Huang J, Tang P, Zhou Y, et al. Mechanisms for U2AF to define 3’ splice sites and regulate alternative splicing in the human genome. Nat Struct Mol Biol. 2014;21(11):997–1005. doi: 10.1038/nsmb.2906. PubMed PMID: 25326705; PubMed Central PMCID: PMCPMC4429597.

27. Cho S, Moon H, Loh TJ, Jang HN, Liu Y, Zhou J, et al. Splicing inhibition of U2AF65 leads to alternative exon skipping. Proc Natl Acad Sci U S A. 2015;112(32):9926–31. doi: 10.1073/pnas.1500639112. PubMed PMID: 26216990; PubMed Central PMCID: PMCPMC4538632.

28. Christopherson JB. The successful use of antimony in bilharziosis administered as intravenous injections of antimonium tartaratum (tartar emetic). Lancet. 1918;192(4958):325-7.

29. Thomas CM, Timson DJ. The Mechanism of Action of Praziquantel: Six Hypotheses. Curr Top Med Chem. 2018;18(18):1575–84. doi: 10.2174/1568026618666181029143214. PubMed PMID: 30370849.

30. Pica-Mattoccia L, Cioli D. Sex- and stage-related sensitivity of *Schistosoma mansoni* to *in vivo* and *in vitro* praziquantel treatment. Int J Parasitol. 2004;34(4):527–33. doi: 10.1016/j.ijpara.2003.12.003. PubMed PMID: 15013742.

31. Sabah AA, Fletcher C, Webbe G, Doenhoff MJ. *Schistosoma mansoni*: chemotherapy of infections of different ages. Exp Parasitol. 1986;61(3):294–303. Epub 1986/06/01. doi: 10.1016/0014-4894(86)90184-0. PubMed PMID: 3086114.

32. Zheng Y, Dong L, Hu C, Zhao B, Yang C, Xia C, et al. Development of chiral praziquantel analogues as potential drug candidates with activity to juvenile *Schistosoma japonicum*. Bioorg Med Chem Lett. 2014;24(17):4223–6. doi: 10.1016/j.bmcl.2014.07.039. PubMed PMID: 25127102.

33. Cioli D, Pica-Mattoccia L. Praziquantel. Parasitol Res. 2003;90 Supp 1:S3-9. Epub 2003/06/18. doi: 10.1007/s00436-002-0751-z. PubMed PMID: 12811543.

34. Alwan SN, Taylor AB, Rhodes J, Tidwell M, McHardy SF, LoVerde PT. Oxamniquine derivatives overcome Praziquantel treatment limitations for Schistosomiasis. PLoS Pathog. 2023;19(7):e1011018. doi: 10.1371/journal.ppat.1011018. PubMed PMID: 37428793; PubMed Central PMCID: PMCPMC10359000.

35. Xu J, Wang JY, Huang P, Liu ZH, Wang YX, Zhang RZ, et al. Schistosomicidal effects of histone acetyltransferase inhibitors against *Schistosoma japonicum* juveniles and adult worms in vitro. PLoS Negl Trop Dis. 2024;18(8):e0012428. doi: 10.1371/journal.pntd.0012428. PubMed PMID: 39159234; PubMed Central PMCID: PMCPMC11361729.

36. Abdul-Ghani R, Loutfy N, Sahn AE, Hassan A. Current chemotherapy arsenal for schistosomiasis mansoni: alternatives and challenges. Parasitology Research. 2009;104(5):955–65.

37. Tevyashova AN, Shtil AA, Olsufyeva EN, Simonova VS, Samusenko AV, Preobrazhenskaya MN. Carminomycin, 14-hydroxycarminomycin and its novel carbohydrate derivatives potently kill human tumor cells and their multidrug resistant variants. J Antibiot (Tokyo). 2004;57(2):143–50. doi: 10.7164/antibiotics.57.143. PubMed PMID: 15112963.

38. Woldemichael GM, Turbyville TJ, Linehan WM, McMahon JB. Carminomycin I is an apoptosis inducer that targets the Golgi complex in clear cell renal carcinoma cells. Cancer Res. 2011;71(1):134–42. doi: 10.1158/0008-5472.CAN-10-0757. PubMed PMID: 21199801; PubMed Central PMCID: PMCPMC3074515.

39. Gause GF, Dudnik YV. Antitumor Antibiotic Carminomycin: Mechanism of Action. In: Hellmann K, Connors TA, editors. Chemotherapy: Cancer Chemotherapy II. Boston, MA: Springer US; 1976. p. 169-73.

40. Basch PF. Cultivation of Schistosoma mansoni in vitro. I. Establishment of cultures from cercariae and development until pairing. J Parasitol. 1981;67(2):179–85. PubMed PMID: 7241277.

41. Oliveira MP, Correa Soares JB, Oliveira MF. Sexual Preferences in Nutrient Utilization Regulate Oxygen Consumption and Reactive Oxygen Species Generation in *Schistosoma mansoni*: Potential Implications for Parasite Redox Biology. PLoS One. 2016;11(7):e0158429. doi: 10.1371/journal.pone.0158429. PubMed PMID: 27380021; PubMed Central PMCID: PMCPMC4933344.

42. Wang J, Collins JJ, 3rd. Identification of new markers for the *Schistosoma mansoni* vitelline lineage. Int J Parasitol. 2016;46(7):405–10. doi: 10.1016/j.ijpara.2016.03.004. PubMed PMID: 27056273; PubMed Central PMCID: PMCPMC4917872.

43. Wang J, Chen R, Collins JJ, 3rd. Systematically improved in vitro culture conditions reveal new insights into the reproductive biology of the human parasite *Schistosoma mansoni*. PLoS Biol. 2019;17(5):e3000254. doi: 10.1371/journal.pbio.3000254. PubMed PMID: 31067225; PubMed Central PMCID: PMCPMC6505934.

44. Collins JJ, 3rd, Hou X, Romanova EV, Lambrus BG, Miller CM, Saberi A, et al. Genome-wide analyses reveal a role for peptide hormones in planarian germline development. PLoS Biol. 2010;8(10):e1000509. doi: 10.1371/journal.pbio.1000509. PubMed PMID: 20967238; PubMed Central PMCID: PMCPMC2953531.

45. Collins JJ, 3rd, Wang B, Lambrus BG, Tharp ME, Iyer H, Newmark PA. Adult somatic stem cells in the human parasite Schistosoma mansoni. Nature. 2013;494(7438):476-9. Epub 2013/02/22. doi: 10.1038/nature11924. PubMed PMID: 23426263; PubMed Central PMCID: PMCPMC3586782.

46. Kim D, Paggi JM, Park C, Bennett C, Salzberg SL. Graph-based genome alignment and genotyping with HISAT2 and HISAT-genotype. Nat Biotechnol. 2019;37(8):907–15. Epub 2019/08/04. doi: 10.1038/s41587-019-0201-4. PubMed PMID: 31375807; PubMed Central PMCID: PMCPMC7605509.

47. Shumate A, Wong B, Pertea G, Pertea M. Improved transcriptome assembly using a hybrid of long and short reads with StringTie. PLoS Comput Biol. 2022;18(6):e1009730. Epub 2022/06/02. doi: 10.1371/journal.pcbi.1009730. PubMed PMID: 35648784; PubMed Central PMCID: PMCPMC9191730.

48. Love MI, Huber W, Anders S. Moderated estimation of fold change and dispersion for RNA-seq data with DESeq2. Genome Biol. 2014;15(12):550. Epub 2014/12/18. doi: 10.1186/s13059-014-0550-8. PubMed PMID: 25516281; PubMed Central PMCID: PMCPMC4302049.

49. Shen S, Park JW, Lu ZX, Lin L, Henry MD, Wu YN, et al. rMATS: robust and flexible detection of differential alternative splicing from replicate RNA-Seq data. Proc Natl Acad Sci U S A. 2014;111(51):E5593–601. Epub 2014/12/07. doi: 10.1073/pnas.1419161111. PubMed PMID: 25480548; PubMed Central PMCID: PMCPMC4280593.

